# Cytokine-induced memory-like responses in endothelial cells link chronic inflammation to vascular disease risk

**DOI:** 10.1101/2025.06.16.659872

**Authors:** Kieu T.T. Le, Nick Keur, Heleen Middelkamp, Thuy Linh Do, Albert van den Berg, Valeria Orlova, Leo A.B. Joosten, Mihai G. Netea, Cisca Wijmenga, Iris Jonkers, Sebo Withoff, Andries D. Van der Meer, Vinod Kumar

## Abstract

Chronic inflammation plays a central role in the progression of both infectious and vascular diseases, yet its impact on endothelial cells (ECs), which form the interface between blood and tissue, remains poorly understood. Given their constant exposure to inflammatory cytokines such as TNF-α and IFN-γ, we set out to investigate how cytokine induced inflammation shapes EC function at the molecular level. Using primary human umbilical vein endothelial cells (HUVECs), we modeled repeated cytokine exposure to simulate a chronically inflamed microenvironment. Transcriptomic and epigenetic profiling revealed that ECs respond to this chronic stimulation with durable transcriptional and chromatin changes. These responses included phenotypes resembling immune cell priming, training, and tolerance, which are commonly associated with innate immune memory, a phenomenon whereby innate immune cells mount altered response following previous stimulation. Although we did not observe classical trained immunity pathways, several genes known to mediate immune training, including *TLR2*, *IL1B*, and *HDAC9*, exhibited persistent activation following TNF-α re-exposure. IFN-γ stimulation uniquely induced sustained expression and chromatin accessibility at MHC class II loci, suggesting cytokine-specific modes of reprogramming. Functionally, re-stimulated ECs exhibited enhanced monocyte adhesion in a 3D vessel-on-chip model, highlighting the relevance of these molecular changes to vascular inflammation. Moreover, the regulatory regions altered by cytokine exposure were enriched for disease-associated SNPs, particularly those linked to COVID-19, sepsis, and cardiovascular disorders. In summary, these findings reveal that repeated exposure to cytokines as seen in chronic inflammation can induce memory-like responses in ECs and suggest that endothelial reprogramming may contribute to vascular dysfunction.

## Introduction

Chronic inflammation is a central feature of many human diseases, including cardiovascular disease, arthritis, and sepsis, where prolonged exposure to inflammatory signals leads to persistent immune activation and tissue dysfunction. Endothelial cells (ECs), which line the blood vessels, are not merely passive barriers but actively regulate immune responses by secreting cytokines such as interleukin-6 (IL-6) and interleukin-8 (IL-8), and by guiding immune cell recruitment to sites of infection or injury^1^ During chronic inflammatory conditions, ECs are repeatedly exposed to proinflammatory cytokines like tumor necrosis factor-alpha (TNF-α) and interferon-gamma (IFN-γ), which are secreted by activated immune cells in response to infections, such as those caused by fungal or gram-positive bacteria.^2^ These cytokines are known to induce and sustain endothelial activation and dysfunction, which are often early steps in the pathogenesis of vascular complications such as cardiovascular disease and sepsis. However, while the acute responses of ECs to inflammatory cytokines have been well characterized, the consequences of repeated cytokine exposure on endothelial molecular states remain poorly understood.

Repeated exposure to microbial components or inflammatory cytokines in innate immune cells such as monocytes and macrophages can lead to long-term metabolic and epigenetic reprogramming, a phenomenon known as trained immunity.^3^ This form of innate immune memory enables enhanced responses to subsequent unrelated pathogens and was initially identified in blood monocytes differentiating into macrophages.^3^ Classical inducers of trained immunity, such as beta-glucan and Bacillus Calmette-Guerin (BCG), were shown to upregulate the production of pro-inflammatory cytokines such as IL-1, TNF-α, and IFN-γ by inducing long-term epigenetic and metabolic changes.^3–6^ Interestingly, more recent studies have demonstrated that cytokines themselves, particularly TNF-α and IFN-γ, can induce similar long-term inflammatory phenotypes in immune cells.^7,8^ Given their central role in many human chronic inflammatory diseases and their persistent presence in the inflammatory microenvironment, these cytokines may drive long-lasting changes in other cell types. Despite not directly recognizing pathogens such as *Candida albicans* and *Streptococcus pneumonia,* ECs respond robustly to endogenous mediators such as IFN-γ and TNF-α. (testetss) These responses include upregulation of adhesion molecules, altered cytokine profiles, and changes in endothelial permeability. Moreover, epigenetic changes in trained monocytes have been linked to changes in vascular endothelial growth factor (VEGF) pathway, endothelin signaling, and the trans-endothelial migration pathways that are also critical to endothelial function and inflammation. ^5^ In addition to their innate immune-like activities, ECs regulate adaptive immunity by expressing T-cell co-stimulatory and immune checkpoint receptors, further reinforcing their active role in immune regulation. ^9^

While trained immunity has traditionally been studied in innate immune cells, emerging evidence suggests that other cell types, including ECs, can exhibit similar immune like phenotypes in response to inflammatory stimuli. Recent studies have shown that ECs can exhibit trained immunity responses to specific stimuli such as lipopolysaccharides (LPS), oxidized LDL (oxLDL) and lysophosphatidylcholine (LPC).^10–12^ Taken together, these observations suggest that repeated cytokine exposure may induce persistent, memory-like responses in endothelial cells, similar to what has been observed in innate immune cells. Exploring this concept may help reveal how endothelial activation contributes to disease persistence in chronic inflammation. In this study, we investigated whether repeated stimulation of ECs with TNF-α and IFN-γ, key cytokines involved in chronic and systemic inflammation, induces stable transcriptomic and epigenetic changes. Using primary human umbilical vein endothelial cells (HUVECs), we characterized the global impact of cytokine exposure on gene expression and chromatin accessibility. We further assessed the functional consequences of these changes using a 3D vessel-on-chip model to mimic physiologically relevant vascular environments. Finally, we analyzed how these cytokine-induced molecular signatures relate to inflammatory processes by integrating them with genetic data from disease-associated traits. Together, this work aims to advance our understanding of how repeated cytokine exposure alters endothelial function and whether such responses reflect inflammatory-memory behavior typically associated with innate immune cells.

## Materials and Methods

### Cell Culture and cytokine stimulation

Primary human umbilical vein endothelial Cells (HUVECs) were used for the study. Single donor HUVECs were purchased (Lonza) and cultured in endothelial cell growth medium (PromoCell, C-22111) supplemented with 1% Penicillin/Streptomycin (10.000U/ml) (Gibco) at 37°C, 5% CO_2_ and saturating humidity. Cells were stored at passage 2, thawed, and seeded for experiments at passage 3. Stimuli were added when cells reached about 70% confluency. For stimulation, HUVECs were stimulated with TNF-α (PeproTECH, 300-01A) or IFN-γ (10 ng/ml) for 4 hours (HIT1). To mimic repeated cytokine exposure, HUVECs were first exposed to cytokine stimulation for 24 hours, followed by washing step and incubation in control medium (EGM-2) for 48 hours before being re-exposed for 4 hours to the same stimuli (HIT2). The responses of cells with repeated exposure (HIT2) were compared to cells exposed only once to stimuli (HIT1), which always took place on the same day the second stimulation was added. Two control conditions were included, the first involved cells that received no stimulation and were cultured only in medium (CTRL1), and the second included cells exposed to a single 24 hours exposure at the same time point as the first stimulation in the repeated condition (CTRL2).

### Flow cytometry

To determine protein expression of adhesion molecules on the cell membrane, the cells were washed with phosphate-buffered saline (PBS), detached using trypsin, washed with PBS, and fixed with Formaldehyde 4% (Thermo). The cell pellets were washed, resuspended in PBS, and stored at 4°C until staining. The cells were divided equally into separate fluorescence-activated cell sorting (FACS) tubes with an ice-cold FACS buffer (PBS supplemented with 5% fetal calf serum, FCS). The cells were stained using the following antibodies: PE-conjugated anti-human E-selectin (CD62E) (Biolegend, 322606), APC-anti human VCAM-1 (CD106) (Biolegend, 305810), FITC-anti-human ICAM-1 (CD54) (Biolegend, 322720) and IgG isotope controls (IgG isotope controls (Biolegend,) for 30 minutes on ice. The cells were washed once and resuspended in the FACS buffer. Samples were analyzed using the Agilent Quanteon system. Multi-color compensation was calibrated using a positive control cell population (TNF-α activated HUVECs). The gating strategy used for FACS analysis is described in the supplementary file (Fig S1).

### RNAseq

Cells were washed in PBS and quickly lysed in the lysis buffer of the MirVanva MagMax RNA isolation kit (Thermofisher) and stored at -20 °C until isolation. RNA was isolated manually according to the manufacturer’s instructions. RNA concentration and integrity were determined using the Bioanalyzer (Agilent D2000). All samples had an RNA integrity number (RIN) score of at least 8. Of these samples 300 ng of RNA were sent to BGI Hongkong for library prep and sequencing on the DNBSEQ platform to achieve paired-end and 150 bp read length.

### ATACseq

To study the impact of stimulation on the cell’s chromatin accessibility, we performed ATAC-seq (Assay for Transposase-Accessible Chromatin with high-throughput sequencing).^13^ In brief, after stimulation, HUVECs were dissociated by trypsin, and 50,000 cells were collected in PBS. A 50 µl of reaction mix containing Transposase enzyme, buffer (Illumina Tagment DNA TDE1 enzyme and buffer kit), and Digitonin (2%) (Promega) was added to the cell pellet as recommended.^14^ The reaction was incubated at 37 °C for 30 min with agitation at 300 rpm, before purifying with a Qiagen MinElute PCR purification kit. The transposed DNA was stored at -20 °C to wait until all samples were ready for PCR amplification. The transposed DNA fragments were amplified into 4-6 cycles (NEBNext High-Fidelity 2X PCR Master mix), before being cleaned again with a Qiagen MinElute PCR purification kit. DNA fragments (100-1000 bp) were selected from the library by bead purifications (Agencourt AMPure XP beads, Fisher Scientific). The quantity and quality of the ATACseq libraries were calculated by Bioanalyzer (Agilent TapeStation), HS D1000. Indexed libraries were pooled and sequenced by BGI on the DNBSEQ platform using a dual index design. Demultiplexed libraries yielded between 46-54 million reads per sample).

### RNA Sequencing Library Processing and Quantification

The obtained raw sequence reads were initially filtered using SOAPnuke. We applied filters which included the removal of sequence adapters, contamination check, and the removal of poor-quality reads (Q ≥ 33). Subsequently, the processed data were evaluated for subsequent quality control using FastQC (version 0.11.9) and MultiQC (version 1.14). The remaining filtered data were then aligned and annotated using the human GENCODE Version 39 (GRCh38.p13) genome using the STAR Aligner (version 2.7.10a) with default parameters. Finally, quantification of reads per gene was performed using STAR’s --quantMode GeneCounts.

### ATAC Sequencing Library Processing and Peak Calling

All samples were processed (alignment and quality check) using the nf-core ATACseq.^15^ In brief, the quality of ATACseq reads was assessed using FASTQC, and adapters were trimmed using TrimGalore. Subsequently, reads were aligned to the human genome (GRCh38.p13) using BWA where the Encode GR38 blacklist was utilized to filter out problematic regions. We then excluded reads mapped to mitochondrial DNA and retained only properly paired reads with high mapping quality (MAPQ score >10, qualified reads) using SAMTools. Next, we removed duplicate reads with Picard, using the MarkDuplicates program and narrow peaks were called using MACS. (parameters: narrow_peak, –paired_end, --deseq2_vst, min_reps_consensus 5, --macs_fdr 1e-5). A peak was deemed genuine if present in a minimum of 5 biological replicates out of the 19 samples. Lastly, HOMER annotatePeaks.pl was utilized to annotate the peaks to known genomic elements. One sample in the CTRL2 condition failed quality control due to extremely low sequencing depth and was excluded from downstream analyses.

### Differential expression analysis

Identification of differential expressed genes (DEGs) and differential open chromatin regions (DORs) were performed using generalized linear models (GLM) in DESeq2 (v1.30.1).^16^ In brief, the DESeq2 algorithm models the gene expression counts using a negative binomial distribution and generates linear models to determine differentially expressed genes. To ensure robust statistical power, genes with read counts of less than 20 and measured in minimal 3 samples were removed before further analysis. To account for inter-individual variability, a paired design was employed, which included each individual’s donor ID as a covariate.

To directly assess transcriptional and chromatin accessibility differences following single and repeated cytokine stimulation, we utilized a contrast-based approach rather than testing interaction terms in a linear model for both RNA-seq and ATAC-seq data. First, we assessed the effects of single cytokine stimulation by comparing HIT1 vs CTRL1, which allows us to explore the immediate transcriptional and chromatin changes induced by cytokine. Next, to explore the response to restimulation, we compared HIT2 vs CTRL2, assessing whether repeated stimulation led to an altered cellular response. To specifically test the effects of cytokine restimulation, we compared HIT2 vs HIT1, identifying genes and open chromatin regions that responded differently upon single and repeated stimulation, independent of the baseline effects. To evaluate potential residuals or memory-like effects we compared CTRL2 vs CTRL1, determining whether changes persisted after initial cytokine stimulation and recovery phase. Lastly, to directly account for residual effects and identify transcriptional dynamics specific to repeated stimulation, we performed a baseline-controlled comparison (HIT2 - CTRL2) vs (HIT1 - CTRL1), this approach allowed us to identify differences in open chromatin regions and gene expression responses between the single and repeated stimulation. This approach captures both genes and regulatory regions that exhibit different dynamics between single and repeated stimulation and are therefore more likely to reflect memory-like behavior.

For ATAC-seq, differentially open chromatin regions (DORs) were identified using the same comparisons as RNA-seq. Additionally, we integrated RNA-seq and ATAC-seq data by mapping DORs to DEGs within a ±10kb window around gene transcription start sites (TSS). The resulting P-values were adjusted across all genes and peaks using the Benjamini-Hochberg method, and adjusted P-values below 0.05 and ± log2FoldChange of 0.5 were considered statistically significant. A complete list of all differentially expressed genes (DEGs), along with their corresponding log2 fold changes and adjusted p-values, is provided as a supplementary file (Table S1)

### Transcriptomic K-means clustering analysis

To discern similar expression patterns across sample groups we employed the ‘degPatterns’ function from the DEGreport R package.^17^ This function calculates the pairwise correlations among all input genes, conducts hierarchical clustering, and trims the generated tree to define clusters/groups with analogous gene expression. The identified clusters were subsequently used as input for functional enrichment analysis. Hierarchical clustering and the creation of heatmaps were executed using the ComplexHeatmap (version 2.14.0) and the pheatmap packages in R (version 1.0.12).

### Functional enrichment analysis

Functional enrichment and Gene Set Enrichment Analysis (GSEA) was conducted to identify pathways and molecular functions associated with DEGs of interest. This analysis was performed using the R packages ClusterProfiler (version 4.6.0) and enrichR (version 3.1), which enables a comprehensive investigation of functional annotations. Additionally, GSEA was used to identify gene sets based on ranked gene expression data, allowing for detection of coordinated changes in an unbiased way and in the absence of strong differential expression. We retrieved gene sets and annotations from several databases including GO terms, KEGG Pathways, and Reactome. To ensure robustness of our results, we applied multiple testing correction using the false discovery rate (FDR), terms and pathways with an FDR corrected p-value < 0.05 were considered significant. Results were visualized using ggplot2 (version 3.4.0), employing bar plots, point heatmap and GSEA enrichment plots to summarize and interpret the enriched pathways.

### Motifs enrichment analysis of chromatin accessibility

To identify enriched motifs in the identified DORs we utilized the R package marge (version 0.0.4). This tool provides an R interface for performing Genomic Analysis using the HOMER (Hypergeometric Optimization of Motif Enrichment) software using tidy data conventions. Motif enrichment analysis was utilized and ran separately on DORs from each pairwise analysis with the flags -size and -mask. The flag -size ensures the use of specific window size around peaks, enhancing the detection of motifs, while the flag -mask removes repetitive elements, reducing noise and improving specificity. Enriched motifs within DORs were determined by comparing the rank differences (based on P-values).

### GWAS SNP analysis of chromatin accessibility

To learn more about the potential functional significance of the identified DORs, we analyzed their overlap with single nucleotide polymorphisms (SNPs) using a permutation-based approach. We extracted all relevant Disease-associated SNPs and annotations from the Genome-Wide Association Study (GWAS) Catalog using all underlying traits from EFO traits list, ’EF_disease’ (EFO:0000408). We then proceeded and tested for enrichment using a permutation-based enrichment test. To ensure robustness and to account for SNP biases, a random set of peaks was sampled from the background of non-differentially open peaks, ensuring that random peaks matched the size and genomic distribution characteristics of the DORs peak set. We determined the overlapping of SNPs with our random samples peak set and the distribution of overlapping SNPs was used to generate a null model (n = 1000). Enrichment significance was determined by comparing the observed overlap of SNPs in the DORs to this null distribution, generating empirical P-values. To approach allowed us to assess whether the disease-associated SNPS were overrepresented in DORs compared to what would be expected by chance, while accounting for confounding factors, such as genomic feature biases. Subsequently, we intersected these SNPs with the GWAS catalog, a comprehensive repository of genetic variants associated with various diseases and traits (Accessed September 2024). We used the ’trackplot’ (version 1.5.1) package to visualize both transcriptomic and epigenetic data, we used the, which enables detailed visualization of RNA-seq and ATAC-seq data as tracks along specific genomic regions.^17^ This tool allowed us to overlay both chromatin accessibility and gene expression patterns and link transcriptomic changes to epigenomic alterations.

### Monocyte adhesion assay on blood-vessel on-chip model Chip fabrication

The microfluidic chip consists of 6 channels, which are one centimeter in length and 500 x 500 µm in width and height (figure 2a-b). The chip was fabricated by conventional polydimethylsiloxane (PDMS)-based soft lithography using a poly (methyl methacrylate) (PMMA, Arkema innovative chemistry) mould. The mould was produced by micromilling (Sherline, model 5410) based on designs in SolidWorks (Dassault Systèmes, France). PDMS (10:1 base:crosslinker ratio, Sylgard 184) was added to the PMMA mould and was left to cure overnight at 65 °C after which it was removed from the mould. 1 mm inlet and outlet holes were created using biopsy punchers (Robbins Instruments). PDMS was spin coated (SPS Spin150) on a glass microscope slide and PDMS was left to cure overnight at 65 °C. The surfaces of both the microfluidic device and the microscope slide were activated by exposing them to air plasma (50 W) for 40 s (Cute, Femto Science, South Korea), after which the microfluidic chip was bonded to the microscope slide. Microfluidic chips were further activated using a 0.22 µm filtered 2 mg/ml Dopamine Hydrochloride (Sigma Aldrich, #H8502) in 10 mM TrisHCl (pH 8.5) solution. This solution was added to the channels and incubated at room temperature for one hour. After one-hour chips were washed 3 times with sterile milliQ.

### Collagen solution

A 5 mg/ml rat tail collagen solution was prepared, using high concentration collagen (Corning life sciences; #354249) which was diluted using a master mix consisting of 10x PBS, 1M NaOH and dH_2_O (always working on ice), while 1/10 of the final volume of 10x PBS was added to the master mix. The amount of 1M NaOH added was calculated using the following formula: (volume collagen) x 0.023. The amount of dH_2_O was calculated and added to the master mix: (Final volume) -(volume collagen)-(volume 10x PBS)-(volume NaOH)= volume dH_2_O. The master mix was mixed thoroughly by pipetting up and down without inducing air bubbles. The collagen was added into a new ice-cold Eppendorf tube using ice-cold pipette tips. Master mix was added to the collagen and mixed gently until mixed properly. Using pH paper, the pH of the solution was checked to be between 6.5 and 7.5. Then the solution was vortexed and spun down to remove air bubbles.

### Viscous finger patterning

Viscous finger patterning inside the microfluidic chips was performed using the published method.^18^ A 200 µl pipet tip cut off to 7 mm was inserted in the outlet of the channel. 10 µl collagen solution was pipetted into the inlet of the channel until a meniscus of collagen formed on the cut pipet tip. The pipet tip was left in the inlet and 2.2 µl of ice-cold PBS was added to the cut pipet tip. This droplet formed a lumen in the collagen inside the channel. After patterning, the microfluidic chip was placed inside an incubator at 37 °C and 5% CO_2_ for 1 hour. After one hour, pipet tips were removed from inlets and outlets and replaced with new 200 µl pipet tips and filled with medium. Chips were incubated overnight at 37 °C and 5% CO_2_.

### Culturing cells in microfluidic chips

HUVEC cells from three donors (donor #SD125; #SD263; #SD126) were thawed at passage and passaged once to obtain enough cells for seeding in microfluidic channels. Cells were detached by trypsin, neutralized, counted by LUNA cell counter and diluted to an end concentration of 5·10^6^ cells/ml. Cells were seeded in the channels using gel loader pipet tips, so as not to remove the present pipet tips from the collagen formed channels. After seeding cells, chips were inverted and incubated at 37 °C and 5% CO_2_ for half an hour to let cells attach to the top of the microfluidic channel. The bottom half of the chip was seeded using a second seeding step. After half an hour of incubation for cell adhesion, all channels were refreshed with 200 µl fresh culture medium (day 0). Chips were placed on a rocking table (10° angle, 30 sec intervals; BenchBlotter™ 2D platform rocker) and after 24 hours (day 1) a monolayer was formed. The endothelial lumens were treated in four different conditions as described previously, including: HIT1, HIT2, CTRL1 and CTRL2 using TNF-α stimulation, while medium was refreshed every 24 hours. Cells were kept in culture for 4 days with specific stimulation conditions and on day 4, the monocyte adhesion assay was performed.

### Monocyte adhesion assay

Monocytes (THP-1) were stained using a cell tracker dye (Invitrogen; C2925), according to recommendations of the manufacturer using a concentration of 2.5 µM and diluted to a concentration of 2.5·10^6^ cells/ml in RPMI 1640 + GlutaMAX medium (Gibco; 61870036). A syringe pump (Harvard Apparatus) was used to pull the monocyte solution through the channels at 2 µl/min. The pump was connected to the outlet of the channel by removing the pipet tip from the channel and adding a 14-gauge blunt needle connected to tygon tubing, which was then connected to the syringe. A brightfield microscope was used to confirm flow. After flow was confirmed, a gel loader tip was used to remove excess medium from the inlet and replaced with the monocyte suspension. Monocytes were perfused through the channel for 10 minutes, after which excess monocyte solution was removed from the inlet pipet and replaced with regular EGM-2 medium. This was pulled through the channel for at least 2 minutes to remove all non-attached monocytes from the channel. After removing all the unattached monocytes in the channel, the pump was stopped and disconnected from the outlet of the channel. The blunt needle was replaced with a yellow pipet tip and chips were used for live imaging.

### Analyzing cell adhesion data

After monocyte adhesion assay, live imaging of microfluidic channels was performed using an EVOS (EVOSTM M5000). Live images were captured on the brightfield channel as well as the GFP channel (monocytes). A pipeline was created in the cell profiler (CellProfiler, Version,4.2.1). The GFP image was first converted to a grayscale image, then primary objects were identified. The thresholding strategy was Global, the thresholding method Otsu. The smoothing filter was adjusted to include as many monocytes as possible. This pipeline was used to identify the number of monocytes in the GFP channel. The number of cells attached to each endothelial lumen was then plotted.

## Results

### Repeated cytokine stimulation model to assess chronic inflammatory and memory-like responses in endothelial cells

To investigate endothelial function and responses to repeated cytokine stimulation, we designed an in vitro stimulation model using two dominant inflammatory cytokines (TNF-α or IFN-γ). This setup was designed to mimic aspects of chronic inflammation and explore whether prior exposure to inflammatory signals influences the transcriptional and functional response to a second stimulation. The cells were initially stimulated for 24 hours, and then washed and allowed to rest for 48 hours before the second stimulation was applied for 4 hours (HIT2). The responses of the repeated stimulated ECs (HIT2) were then compared with ECs that received a single 4-hour stimulation (HIT1). Two control conditions were included: ECs that remained unstimulated throughout (CTRL1), and ECs that were stimulated for 24 hours and then rested for 48 hours without a second stimulation (CTRL2). These conditions were used as controls to assess the impact of the culture’s duration and remnant effects of the first stimulation (Fig. 1A).

**Figure 1:**
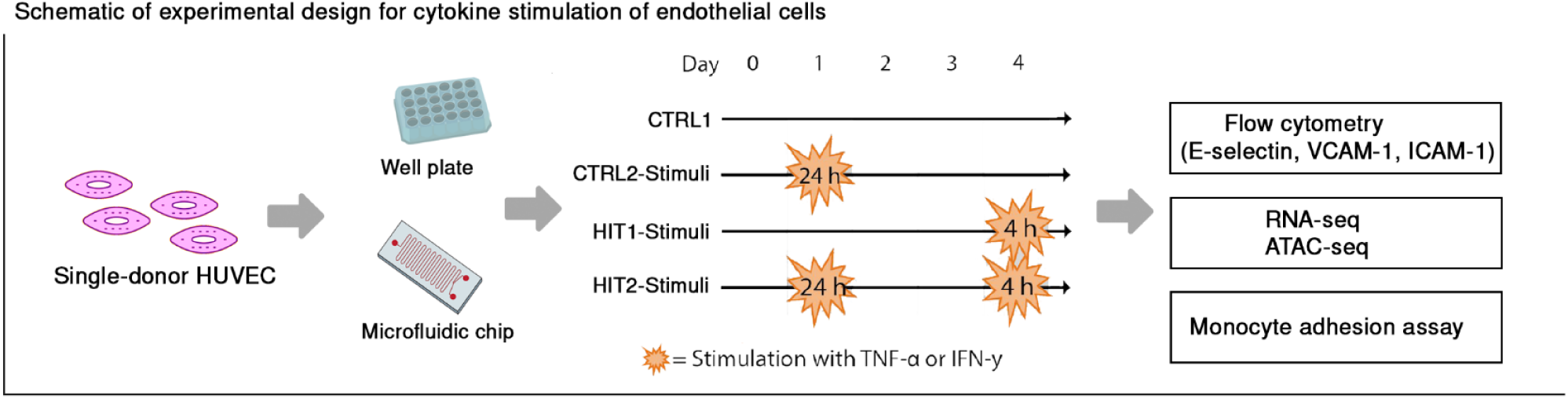
Schematic of experimental design. Endothelial cells were stimulated with either EGM-2, serving as a control (CTRL1), or with a single 24-hour stimulation on day 1 (CTRL2), serving as a control for the second hit of stimulation. To study cellular responses upon repetitive stimulation and contrast it to the first response, ECs were stimulated freshly stimulated with either TNF-α or IFN-γ (HIT1) or received a second stimulation (HIT2) on day 4. On day 2, the stimuli were washed away, and the cells were seeded with fresh medium.

### Global transcriptome changes in endothelial cells after TNF-α or IFN-γ re-exposure

To investigate the global transcriptional responses of ECs to single and repeated cytokine stimulation, we stimulated ECs from five individuals and conducted RNA sequencing of the harvested cells, followed by differential expression analysis. Overall, cytokine stimulation by either TNF-α or IFN-γ induced major transcriptional changes compared to transcriptional changes due to inter-individual differences (Fig. 2A, 2G). For both TNF-α and IFN-γ, more DEGs were identified following a single stimulation than after the repeated stimulation (Fig. 2D, 2J, S2A-D). Upon repeated exposure, the cells activated many genes that were shared with the single exposure, while less new genes were induced (Fig 2C, 2I). The proportion of newly activated DEGs responding to repeated stimulation accounted for 11,3% and 14,5% for TNF-α and IFN-γ, respectively. In addition to a large portion of genes that were uniquely activated following either TNF-α or IFN-γ treatment, indicating cytokine-specific effects on ECs, 280 genes responded commonly to all stimulation conditions (Fig. S3G).

**Figure 2:**
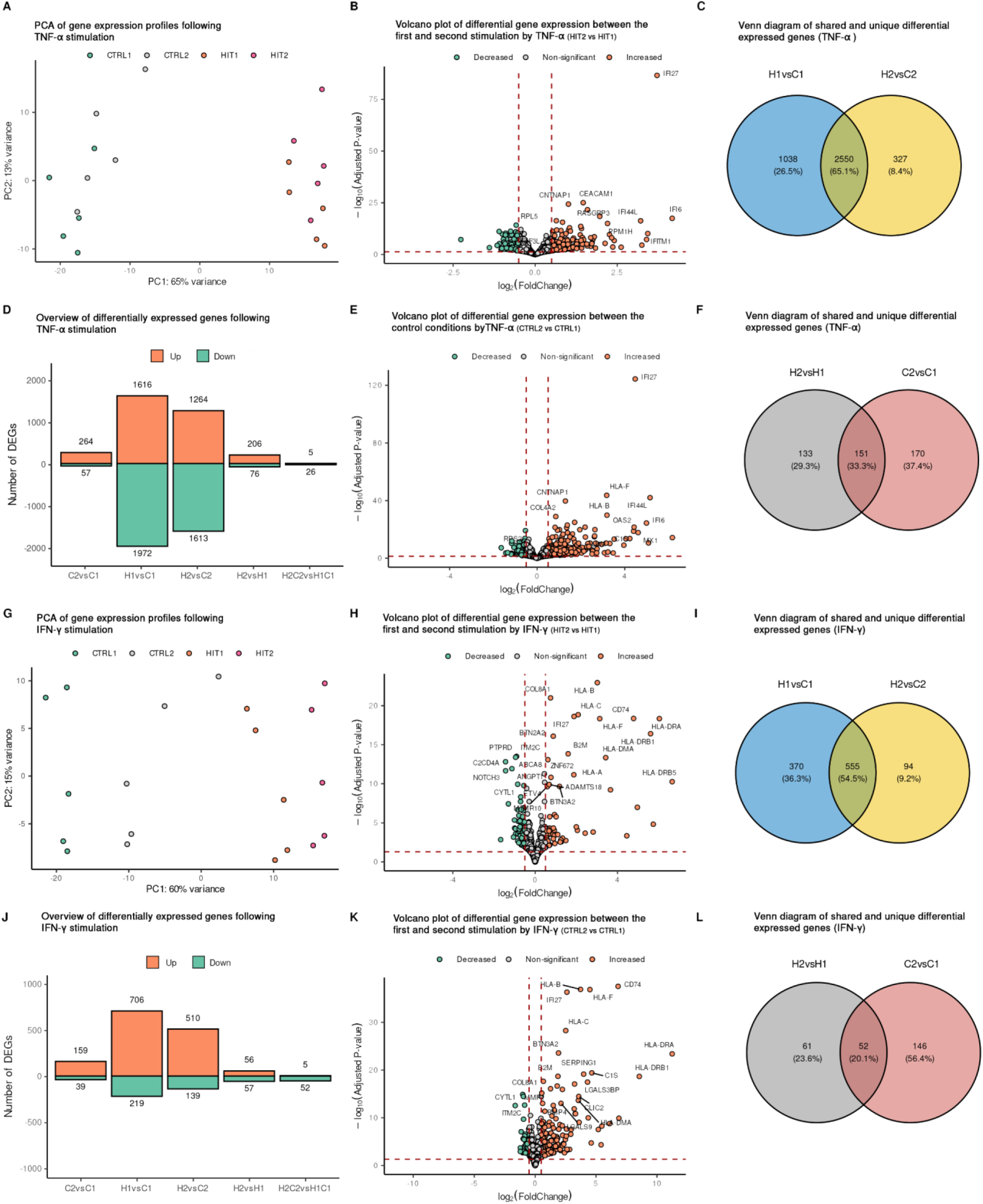
Transcriptome of ECs to various stimulation conditions. ECs obtained from five donors were stimulated as described in the methods section, and RNA was harvested for RNA-seq analysis. PCA plots depict the behavior of each sample upon TNF-α stimulation **(A)** and IFN-γ stimulation (**G**). Each color represents one of the four stimulation conditions. Volcano plots showing the DEGs between HIT2 vs HIT1 upon (**B)** TNF-α, (**H**) IFN-γ. Volcano plots showing the DEGs between CTRL2 vs CTRL1 upon (**E**) TNF-α, (**K**) IFN-γ. Bar plots showing the total number of DEGs in each pair-wise comparison upon TNF-α (**D**), IFN-γ stimulation (**J**). DEGs were filtered based on FDR adjusted p-value (adjusted p-value ≤ 0.05 and log2FC ≥ 0.5. Venn diagrams illustrating the overlap of DEGs between the first and second stimulations for TNF-α **(C)** and IFN-γ **(I)**. Venn diagrams show the overlap between DEGs identified in HIT1 vs HIT2 and CTRL1 vs CTRL2 comparisons for TNF-α **(F)** and IFN-γ **(L)**, indicating genes involved in long term activation and immune memory-like response.

Next, we assessed the molecular consequences of repeated cytokine stimulation by comparing it to a single stimulation. For the TNF-α stimulation, we identified 282 DEGs when comparing single stimulated and restimulated ECs (HIT2 vs HIT1) (Fig. 2B). Among these, 53% (151 genes) remained elevated even after a 48-hour rest following the first TNF-α stimulation, which suggests a persistent transcriptional state change (Fig. 2E). In contrast, 133 DEGs were altered upon re-stimulation and not affected by residual effects of the first stimulation. This subset may include genes that are newly induced or repressed in response to repeated TNF-α stimulation, potentially representing a secondary or memory-like transcriptional response (Fig. 2F). Regarding the IFN-γ stimulation, we identified 113 DEGs when comparing single stimulated cells and restimulated (HIT2 vs HIT1) (Fig. 2H, 2J). Among these, 46% (52 genes) remained elevated after a single stimulation with IFN-γ, while the remaining 54% (61 genes) were altered upon restimulation (Fig. 2K, 2L). In addition to comparing single-stimulation cells to restimulated cells (HIT2 vs HIT1), we assessed the effects of restimulation relative to baseline conditions (HIT2 - CTRL2 vs HIT1 - CTRL1). This baseline-adjusted approach accounts for persistent differences, ensuring a more precise interpretation of the response magnitude and direction of the restimulation. For the TNF-α stimulations we identified a total of 31 DEGs, whereas for IFN-γ we identified 57 genes, indicating that the transcriptional dynamics changed between single and repeated stimulations. Specifically, repeated IFN-γ stimulation of ECs resulted in 5 genes showing an increased response (upregulated) and 26 genes showing a decreased response (downregulated). Similarly, repeated TNF-α stimulation led to 5 genes being upregulated and 52 genes being downregulated. These findings indicate that repeated stimulation alters the magnitude of transcriptional responses, either enhancing or suppressing gene expression in restimulated ECs. Interestingly, TNF-α and IFN-γ showed different transcriptional signatures. The IFN-γ response was skewed towards gene upregulation, whereas TNF-α resulted in slightly more down regulated genes. This directional asymmetry likely reflects the distinct signaling pathways and gene regulatory programs activated by each cytokine.

To explore common molecular properties of TNF-α and IFN-γ in inducing transcriptional reprogramming and immune memory, we intersected four DE gene sets for both cytokine stimulations (Fig. S3). We identified 13 DEGs between the first and second stimulations of both cytokines which also differed between the control conditions (Fig. S3H). These findings indicate that these genes are activated early and remain activated throughout the different stimulations and resting periods. These genes are *OLFML3*, *PCDH12*, *ADTRP*, *BTN3A2*, *HLA-F*, *HLA-A*, *HLA-C*, *HLA-B*, *IFITM3*, *IFI27*, *B2M*, *LGALS3BP,* and *TYMP,* most of which are known to be involved in the interferon signaling pathway. In addition, no common genes were identified upon repeated exposure to TNF-α and IFN-γ.

### Transcriptome changes in endothelial cells after TNF-α or IFN-γ re-exposure indicate both “trained”, and “primed” genes

To characterize transcriptional expression patterns in response to single and repeated stimulations, we conducted unsupervised clustering of DEGs identified in at least one condition involving cytokine exposure. This included genes responsive to single stimulation, repeated stimulation, or those showing altered expression upon repeated stimulation. For TNF-α stimulation, clustering analysis revealed eight distinct clusters based on transcriptional dynamics (Fig. 3A-B). Functional enrichment was then performed for each cluster to characterize their biological relevance (Fig. 3C-F). Cluster 1 displays a progressive and steady increase in gene expression across stimulated conditions. Pathway enrichment analysis revealed associations with antigen presentation (*HLA-A, HLA-B, TLR1*), Interferon signaling (*RSAD2, ISG15, OAS2*) and the broader immune response pathway, suggesting a role in maintaining immune function. Cluster 2 is characterized by increased expression following cytokine stimulation, followed by a decrease that did not return to baseline expression levels. This pattern of sustained transcriptional elevation upon cytokine stimulation is consistent with primed and trained immunity.^19^ This cluster includes genes such as *TLR2, IL1B,* and *HDAC9*, which are known to be involved in inducing trained immunity in monocytes.^4,20,21^ Other enriched pathways include extracellular matrix organization, nicotinate metabolism (*BST1, PARP8, PARP10*) and cytokine−mediated signaling pathway (*CD74, IL6, IL12, IL34*). Cluster 3 includes a large set of 1241 genes that are robustly induced upon TNF-α stimulation. Expression levels show an initial increase in expression followed by returning to baseline in the resting condition (CTRL2) and are reinduced upon repeated stimulation, reaching similar levels or having a dampened response. This pattern reflects cytokine-responsive with variability upon repeated stimulation. Enriched pathways are marked by strong associations with inflammatory signaling pathways, including TNF signaling, TLR cascades, chemotaxis and tissue morphogenesis. Cluster 4 displays no immediate effect after cytokine stimulation but displays differences after the cells have rested (delayed response). The genes in this cluster are enriched in pathways related to extracellular matrix (ECM) and ribosome biogenesis.

**Figure 3:**
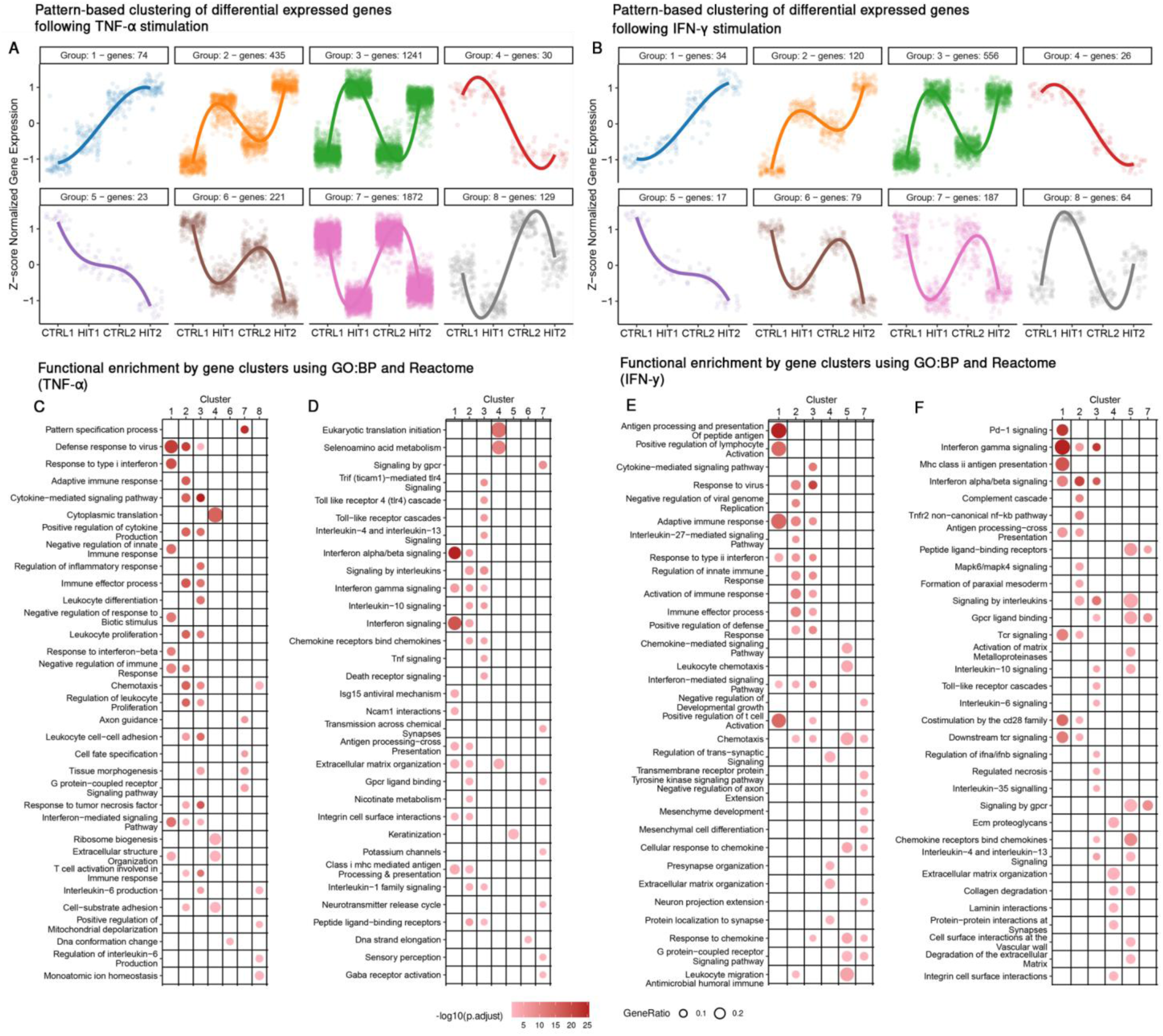
Clustering of RNA expression reveals distinct gene expression patterns associated with immune behaviors (trained, primed and tolerance). Clusters of gene expression patterns upon TNF-α stimulation **(A)**, IFN-γ stimulation **(B)**. The y-axis represents the z-score of the pairwise correlations of genes across all stimulation conditions for each cytokine stimulation, while the x-axis represents the stimulation conditions. Pathway enrichment analysis for DEGs in each cluster is presented for TNF-α using GO biological process terms **(C)** and Reactome pathways **(D),** and for the IFN-γ stimulation using GO biological process terms **(E)** and Reactome pathways **(F)**. The size of each dot represents the gene ratio, while the color represents the adjusted p-value. Clusters with no significant enrichment results are not shown.

Cluster 5 is characterized by a steady decline in gene expression. Functional enrichment analysis revealed that this cluster is associated with keratinization (*KRT7, KRT19* and *MMP28*). This cluster likely reflects a transcriptional program associated with tissue remodeling. Cluster 6 displayed a progressive decrease in expression upon cytokine stimulation, with the lowest levels observed in the repeated stimulation (HIT2). This suggests active suppression or downregulation in response to persistent inflammatory stimulation. Functional enrichment analysis revealed pathways related to DNA strand elongation and DNA conformation change, pointing towards repression of genes involved in DNA replication, repair and chromatin remodeling. Cluster 7 depicts a pattern in which genes are decreased during the first stimulation, return close to baseline levels during the resting phase, and decreased again during the second stimulation, suggesting that these genes are highly regulated during stimulation. The genes in this cluster are enriched in pathways such as signaling by GPCR and tissue morphogenesis suggesting that cells may temporarily halt these processes during cytokine stimulation, while recovering during the resting phase. Cluster 8 includes genes whose expression mildly decreased upon cytokine stimulation, but during the resting phase expression increased above baseline levels, while decreasing again upon restimulation. Despite these minimal changes, pathway enrichment analysis revealed significant associations with biological processes such as chemotaxis (*IL-16, IL17RA*) and mitochondrial depolarization.

Similarly, for the IFN-γ stimulation, we identified 8 pattern-based clusters (Fig. 3B). Interestingly, the global transcriptional dynamics observed between the IFN-γ and TNF-α stimulation groups were similar, which was particularly evident in terms of the observed patterns, such as priming, tolerance, and feedback mechanisms. The observed clusters display comparable dynamics across both cytokines, characterized by pathways involved in immune signaling, antigen presenting, cell cycles regulation and metabolic processes (Fig. 3E-F). However, several distinct features were identified across the TNF-α and IFN-γ stimulation. For instance, while both cytokines strongly activate immune response pathways, IFN-γ clusters display a more pronounced enrichment of interferon-related pathways, such as type II interferon signaling, antiviral response, and antigen processing and presentation. In contrast, stimulations with TNF-α showed a stronger enrichment in pathways related to cellular structural organization and cell division, suggesting a stronger role in driving proliferation and structural adaptation. Furthermore, genes associated with MHC class I antigen presentation were predominantly observed following TNF-α stimulation, while IFN-γ stimulation induced more MHC class II associated genes, highlighting the cytokine-specific antigen presenting programs. Notably cluster 8 was the only cluster with divergent dynamics.

Overall, both cytokines share similar transcriptional dynamics at the global level. Notably, Cluster 2, which displays a primed expression pattern across both cytokine stimulations, stands out. This pattern may reflect a form of transcriptional persistence or primed state in ECs, potentially enabling an enhanced response to recurrent inflammatory stimulation.

### Targeted functional enrichment analysis reveals endothelial and metabolic adaptations

While clustering of gene expression patterns across all conditions provided a global view, we next focused on more targeted comparisons to assess the impact of cytokine stimulation on specific endothelial programs and metabolic adaptations. To assess whether cytokine stimulation activates known endothelial cell transcriptional programs, we performed GSEA using curated endothelial-specific gene sets based on DEGs from directly stimulated comparisons (HIT1 vs CTRL1 and HIT2 vs CTRL2) and used DEGs from the post-stimulation comparison (CTRL2 vs CTRL1) and baseline-controlled comparison (HIT2 - CTRL2 vs HIT1 - CTRL1) to assess metabolic pathways. Analysis of endothelial-specific pathways revealed consistent upregulation of gene sets related to leukocyte adhesion, endothelial activation and interferon signaling across both cytokine stimulations (Fig. S4). Notably, TNF-α stimulation resulted in enrichment for gene sets related to vascular remodeling and endothelial responses, supporting its broader impact on endothelial function. Similarly, analysis of metabolic pathways revealed a modest but consistent shift in endothelial cell metabolism under inflammatory stress, including suppression of glycolysis-related pathways and modest upregulation of nicotinate and nicotinamide metabolism following cytokine stimulation (Fig. S5). These observed metabolic shifts may reflect adaptations in energy production and redox balance under conditions of sustained inflammatory signaling.

### Changes in chromatin accessibility landscape after TNF-α or IFN-γ re-exposure

Part of the mechanism underlying memory-like responses or trained immunity in other cell types is explained by chromatin modifications. To investigate whether similar chromatin modifications occur in ECs in response to repeated cytokine stimulation, we conducted ATAC-sequencing of ECs from the five donors (Fig. 1A). Peak calling was performed on each individual sample, and a consensus peak set was generated for each cytokine stimulation (TNF-α, IFN-γ) independently to enhance reliability and reduce background noise. PCA plots illustrated inter-individual differences (PC1) together with the stimulation effect (PC2) (Fig. S6A, S6E). In concordance with the transcriptome data, one biological replicate strongly varied from the other replicates, regardless of the stimulation.

Upon single stimulation with TNF-α (HIT1 vs CTRL1), we identified 12679 differential open regions (DORs) (Fig. 4A-B), enriched for GO terms including TLR 2,3,4,5 signaling, LPS tolerance and cell activation (Fig. S7A). Repeated exposure led to the identification of 6153 DORs (HIT2 vs CTRL2) (Fig. 4A, 4C) with similar enrichment profiles (Fig. S7B). To assess persistent chromatin accessibility, we analyzed the overlap between single and repeated stimulation and identified 5265 shared DORs representing 42% of the DORs identified in the single stimulation and 86% the repeated stimulation (Fig. S6A-B). Signal intensity plots demonstrated consistent accessibility across these shared regions, while unique DORs showed condition-specific opening or closing (Fig. 4G). Shared DORs were located at the promoter-TSS of genes including *TLR6, IL1R1, IL1B, IL33, NOS3, NFKBIZ*, as well as at the transcription termination sites (TTS) of genes such as *CCL2, SELE, CX3CL1* and *IL8*. These shared DORs highlight conserved chromatin accessibility patterns in response to chronic TNF-α stimulation.

**Figure 4:**
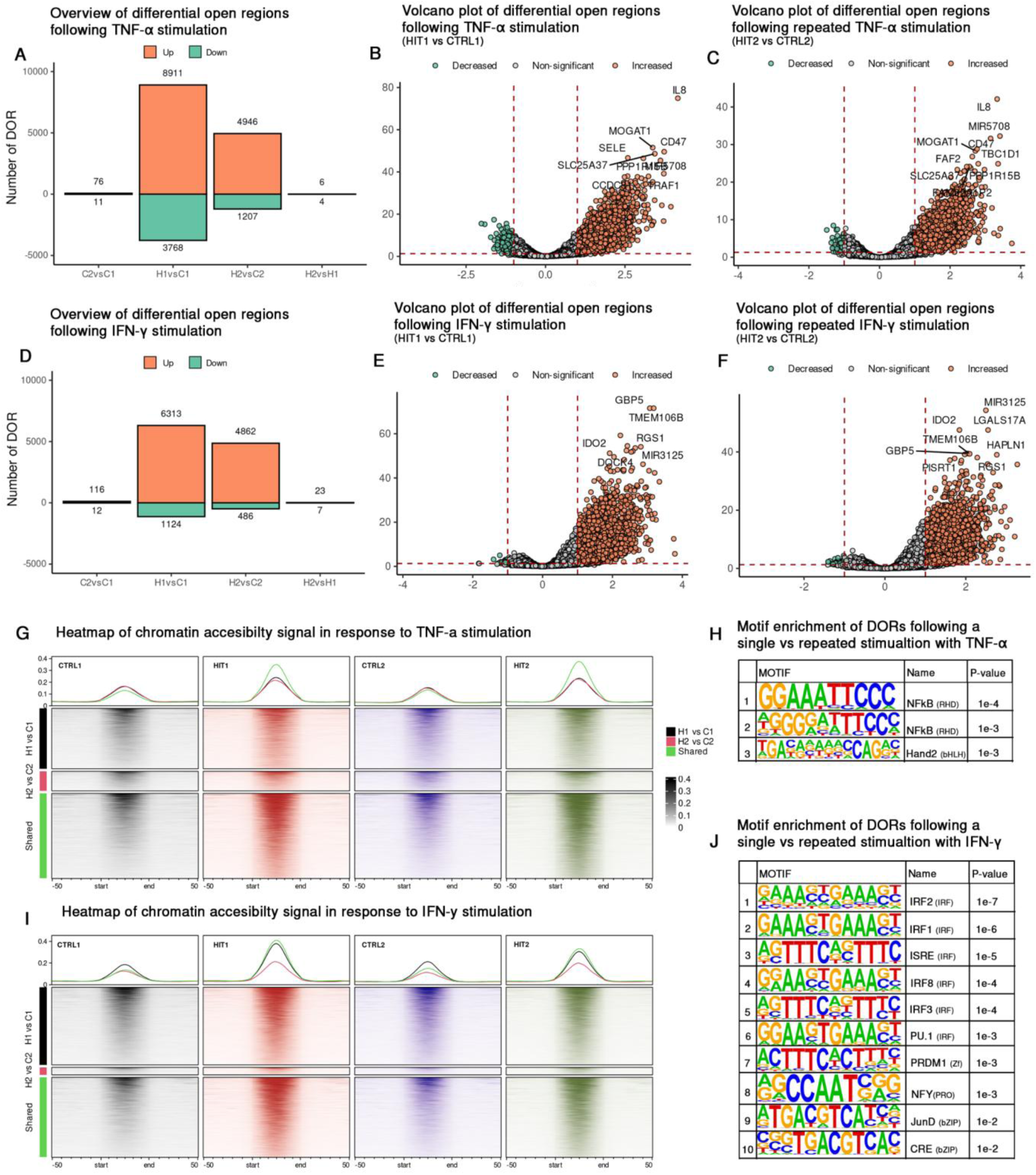
Shared and distinct opening areas of chromatin in EC upon repetitive exposure to TNF-α or IFN-γ. Bar plots showing the total number of DORs for pairwise comparisons identified across pairwise comparisons following TNF-α stimulation **(A)** and IFN-γ stimulation. **(D).** Volcano plots display the total number of DORs detected in ECs for HIT1 vs CTRL1 upon TNF-α **(B)**; HIT2 vs CTRL2 upon TNF-α **(C)**, HIT1 vs CTRL1 upon IFN-γ **(E)**; HIT2 vs CTRL2 upon IFN-γ stimulation **(F)**. Heatmaps illustrate shared and unique chromatin accessibility signals DORs across stimulation conditions, for TNF-α **(G)** and for IFN-γ **(I).** Motif enrichment analysis highlights transcription factor motifs detected in DORs between HIT2 vs HIT1 stimulation upon TNF-α **(H)** and IFN-γ **(J)**.

For IFN-γ stimulation, we identified 7437 DORs following single stimulation (HIT1 vs CTRL1) (Fig. 4D-E), enriched for chemokine production, cell adhesion to the vasculature and cellular response to laminar shear stress (Fig. S7C). Additionally, when examining DORs from repeated stimulation (HIT2 vs CTRL2), 5348 DORs were identified (Fig. 4F), with similar enrichment and additional enrichment for type I interferon signaling, regulation of T-cell, lymphocyte migration, relaxation of cardiac muscle and regulation of IL-1 alpha (Fig. S7D). As with TNF-α, we assessed shared and unique DORs for IFN-γ stimulation (Fig. 4I). We identified 4348 shared DORs, accounting for 58% and 81% of total DORs in single and repeated stimulation conditions, respectively (Fig. S8C-D). Notably, these shared DORs are distributed across the promoter-TSS of 145 genes, including. *ISG15, NLRC5, CASP8, CD74, PIPOX, DDAH1* and *IDO1*, as well as the TTS region of 50 genes.

To determine whether the cytokine-induced chromatin changes persist after stimulation and rest, we compared unstimulated ECs with those that had been exposed to a single 24-hour stimulation and rested for 48 hours (CTRL2 vs CTRL1). For TNF-α, we identified 87 DORs, with 76 upregulated and located at promoters of genes such as *RSAD2, ADAMTS1, ADAM28, SOCS2*. For IFN-γ, we identified 128 DORs, of which 116 were upregulated and therefore still accessible in CTRL2. These DORs were located at promoters of genes including *CD74, IFI27, HLA-DMA, CASP8* and *NOX4*, which are associated with antigen presentation, inflammation and endothelial dysfunction. These persistent chromatin changes may reflect either incomplete resolution of the inflammatory response or priming of regulatory elements for rapid reactivation upon subsequent stimulation.

When we directly compared single and repeated stimulation (HIT2 vs HIT1), we identified 10 and 30 DORs affected by TNF-α and IFN-γ, respectively (Fig. 4A, 4D). For TNF-α, these included promoter regions at *SELE* and *NOD2*, whereas for IFN-γ, promoter DORs were observed at *CD74* and several HLA genes, such as *HLA-DOA, HLA-DMA*, and *HLA-DPB1*. Notably, *CD74* and *HLA-DMA* were among the DORs with persistent accessibility, suggesting that these regions remain epigenetically responsive. Moreover, no DORs were shared between these two cytokine-induced conditions, suggesting a cytokine-specific effect on chromatin regulation during persistent inflammation. In addition to directly comparing single and repeated stimulation, we assessed chromatin accessibility changes while controlling for baseline differences (HIT2 - CTRL2 vs HIT1 - CTRL1). Notably, no significant DORs were identified for either cytokine, suggesting that the chromatin landscape is not dramatically altered upon repeated stimulation, but consistent with those induced in the single hit. Although the number of DORs identified between the single and repeated stimulation conditions was small, the genes associated with these DORs suggest distinct functional pathways for each cytokine. For example, *SELE* and *NOD2* are linked to endothelial activation and immune responses and are considered hallmarks of TNF-α-mediated inflammation. In contrast, the IFN-γ specific DORs highlight pathways involved in antigen processing and presentation via MHC class II, IFN-γ mediated signaling, and cell-cell adhesion. These pathways such as cell-cell adhesion point to the role of interactions between monocytes and endothelial cells during chronic inflammation.

To further understand the differences in cellular responses between single and repeated stimulations, we investigated the binding of different transcription factors within these DORs. We found several transcription factors that were specifically enriched within DORs. Among them, Hand2 is predicted to bind uniquely within DORs that open during the second stimulation of TNF-α (Fig. 4H), whereas four motifs: NFY, CRE, Smad3, and JunD are found to be present within DORs that open upon re-exposure to IFN-γ (Fig. 4I).

### Blood-vessel-on-chip experiments validate the impact of re-exposing endothelial cells to TNF-α on monocyte adhesion

Pathway enrichment of DEGs of the single and repeated stimulations with either TNF-α or IFN-γ indicated changes in the expression of cell adhesion molecules (Fig. 3C-F, S9). However, while repetitive exposure to TNF-α increases the protein expression level of some adhesion molecules, IFN-γ stimulation did not significantly alter the expression of these markers. Given that the transcriptome and epigenetic data suggest inter-individual differences, we examined the expression of these adhesion molecules in cells from individual donors upon TNF-α exposure. Repeated stimulation increased, decreased or did not affect these markers in a donor dependent manner (Fig. 5A). Since changes in adhesion molecules may affect endothelial interactions with immune cells, we employed a blood-vessel on chip model to functionally investigate the impact of repeated TNF-α exposure on endothelial-monocyte adhesion. A PDMS soft-lithography chip was fabricated, and a rat tail collagen type-1 hydrogel was used to create a soft extracellular matrix whereas a round 3D lumen was made inside the collagen using viscous-finger patterning. ECs from three donors (SD125, SD126 and SD263), representing three distinct patterns of expression of adhesion molecules were separately seeded into the lumen. Monocytes were pre-stained with GFP-lipid stain and subsequently used for the adhesion assay. Upon exposing ECs to monocytes, we observed a greater number of monocytes adhering to the endothelial lumen that had been repeatedly exposed to TNF-α compared to those exposed only once (Fig. 5B-C). This suggests that re-exposure of the endothelium to TNF-α alters the functional interaction between endothelial and immune cells. As expected, we observed inter-individual variability among ECs in their capacity to adhere monocytes, suggesting that other factors such as individual genetics may influence the memory-like response of ECs.

**Figure 5:**
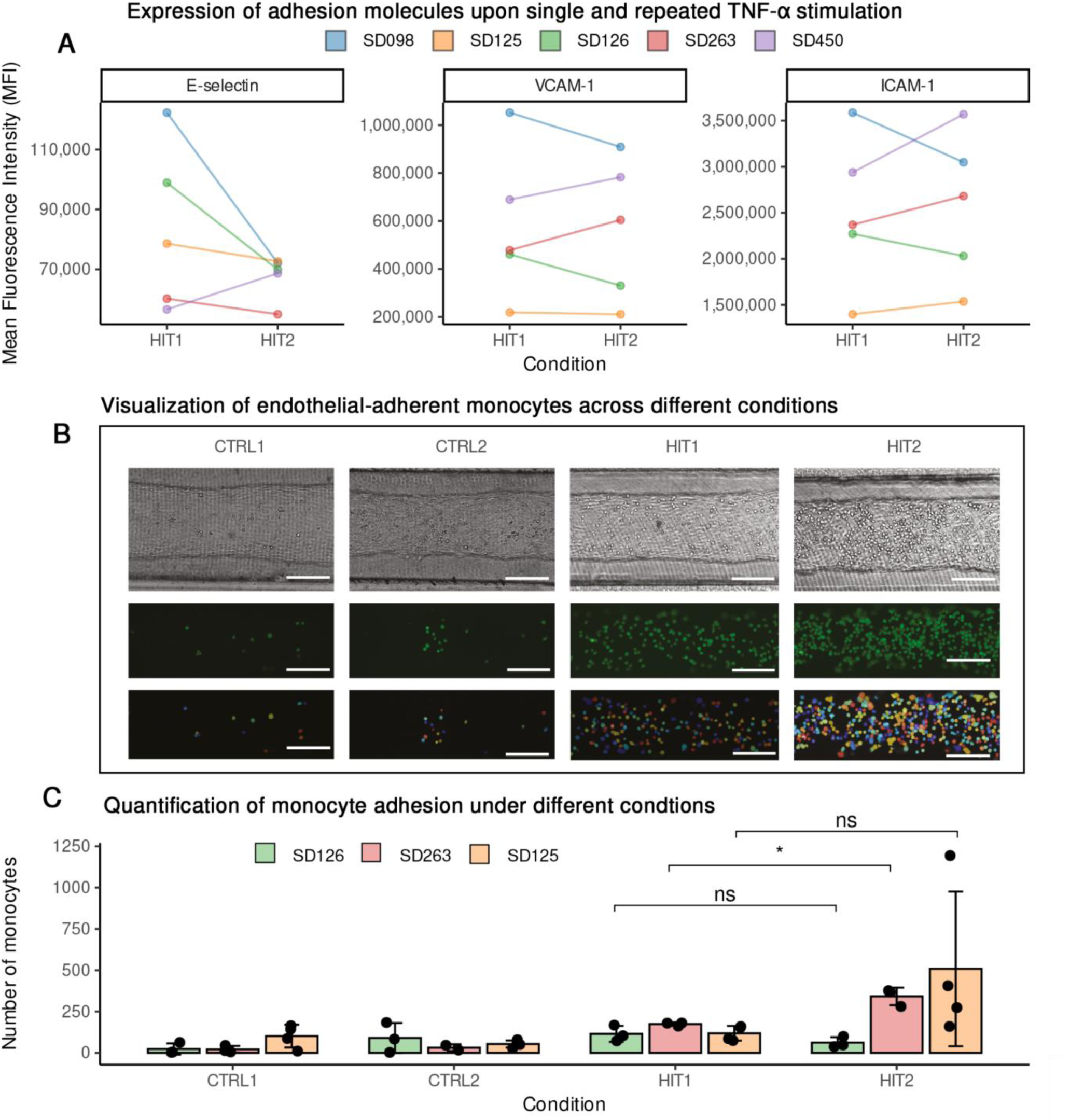
Monocyte adhesion assay on blood-vessel on chip demonstrates the impact of repetitive TNF-α exposure on monocyte-endothelial adhesion. Expression of adhesion markers, measured by flow cytometry, 4 hours after the latest stimulation on primary ECs obtained from five different donors (n=5). Each color indicates cells of the same donor **(A).** The different functional responses of one donor to the different types of stimuli indicate a training effect of the endothelial cells **(B)**. Monocyte adhesion was assessed in three donors with varying responses: a low responding donor showed no increase in monocytes adhesion, whereas medium and high responding donors exhibited and an increase in monocyte adhesion following repeated stimulation. **(C)** All scale bars represent 150 μm.

### Persistent-inflammation-induced chromatin DORs in endothelial cells harbor GWAS-associated variants

To further explore the role of chromatin accessibility dynamics in ECs under chronic inflammatory conditions, we categorized the DORs into two peak-sets for both TNF-α and IFN-γ stimulation: ‘Shared DORs’, which represent conserved chromatin features across single and repeated stimulations, and ‘Dynamic DORs’, which capture differences in chromatin accessibility between the single and repeated stimulation (e.g., difference in responsiveness). Across both cytokine stimulations, DORs in both peak-sets overlapped with GWAS-SNPs linked to diseases such as hypertension, and COVID-19. Among these GWAS-SNPs, 20-30% were observed in gene promoter regions, whereas the others are located in intron and intergenic regions. Other notable GWAS-SNPs are reported to be associated with metabolite levels as well as in infectious diseases. Interestingly, SNPs within DORs that appear upon IFN-γ stimulation are more often associated with immune diseases and immune phenotypes such as systemic lupus erythematosus (SLE) (Fig. 6C), COVID-19 (Fig. 6D), autoimmune diseases (Fig. 6E), and response to vaccines (Fig. 6F). Many of these SNPs overlap with chromatin regions linked to HLA genes, which are critical for antigen presentation and immune regulation. We also observed an SNP rs3087456 (*CIITA* G-286A), that impacts mortality among patients with septic shock.^22^ and that overlaps with the chromatin region that opens strongly after the second hit with IFN-γ (Fig. 6G). *CIITA* is a transcriptional co-activator essential in HLA genes expression.

**Figure 6:**
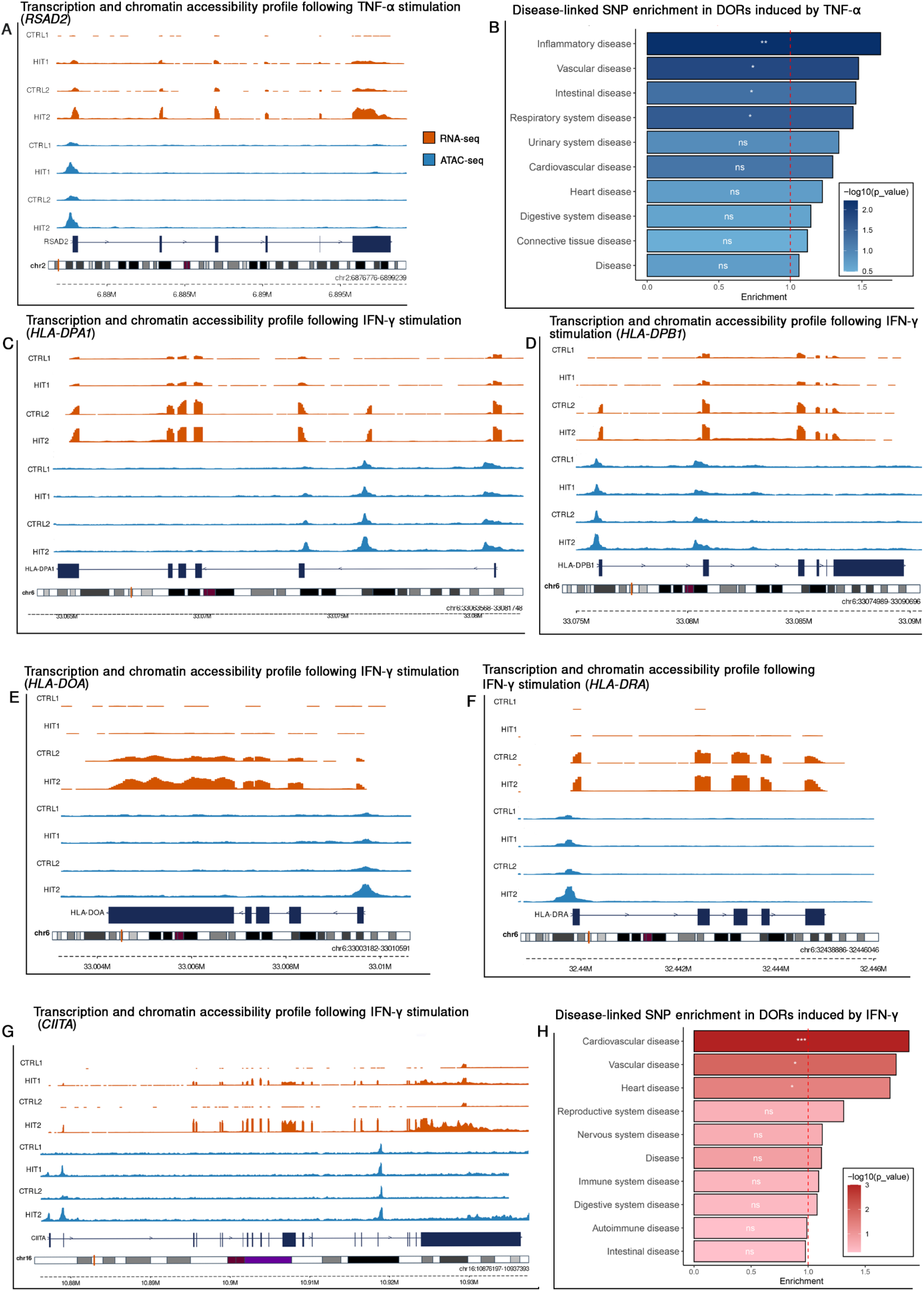
Chromatin accessibility and transcriptional profiles of trained endothelial cells and SNP enrichment in disease-associated traits following TNF-α and IFN-γ stimulation. Transcriptional and chromatin accessibility profile of *RSAD2* (rs75460609) following TNF-α stimulation **(A).** Transcriptional and chromatin accessibility profile following IFN-γ stimulation, *HLA-DPA1* (rs381218) **(C)**, *HLA-DPB1* (rs2071351) **(D)**, *HLA-DOA* (rs381218) **(E)**, *HLA-DRA* (rs2395179) **(F)** and *CIITA (*rs3087456*) (G)*. Orange tracks represent RNA-seq data, and blue tracks represent ATAC-seq data. Genetic profiles for transcriptomic and chromatin signals are scaled for each locus. SNP enrichment analysis of DORs across disease-associated traits is shown for TNF-α stimulation **(B),** and IFN-γ **(H)**. P-values are displayed inside the bars, the red dashed line indicates the significance threshold (p = 0.05), and the bar color represents the enrichment score.

Next, we investigated whether there was significant enrichment of disease-associated GWAS-SNPS in the identified DORs using a permutation-based approach. We separately assessed Shared and Dynamic DORs for both TNF-α and IFN-γ stimulations, resulting in four distinct peak sets. For each set, we generated a null distribution (n=1000) by randomly selecting an equal number of non-differentially open chromatin regions and tested whether the number of DORs with at least one disease associated GWAS-SNP was greater than expected by chance. This analysis revealed that significant enrichment was observed mostly in the peak sets with dynamic DORs (p < 0.05). Dynamic DORs induced by TNF-α stimulation showed statistically significant enrichment with inflammatory diseases, vascular diseases, and intestinal diseases and respiratory system disorders (Fig. 6B). For IFN-γ, dynamic DORs were enriched for cardiovascular, vascular and heart diseases (Fig. 6H). In contrast, shared DORs for both cytokines showed weak or no enrichment across most disease categories. These findings reveal that stimulus-specific, dynamic changes in chromatin accessibility are more likely to intersect with disease-associated genetic variants and may play a role in linking inflammatory history to disease-relevant regulatory elements.

## Discussion

Our study reveals that chronic exposure to proinflammatory cytokines can induce persistent, memory-like responses in human endothelial cells (ECs), providing new insight into how vascular tissue may be reshaped by inflammation over time. By mimicking a chronically inflamed microenvironment using repeated TNF-α and IFN-γ stimulation in primary HUVECs, we demonstrated that ECs undergo transcriptional and epigenetic changes with functional consequences relevant to vascular disease. These memory-like states, although lacking the canonical features of trained immunity seen in monocytes, nevertheless exhibit persistent reprogramming and overlap with genomic regions associated with immune and vascular disorders linking inflammatory history to long-term disease risk.

ECs, while not classical immune cells, express several pattern recognition receptors and exhibit immunomodulatory functions, justifying their characterization as conditional innate immune cells.^23^ Previous studies have shown that ECs can exhibit long-term phenotypic shifts in response to stimuli such as LPS and oxLDL.^10,11^ Extending these findings, we show that repetitive exposure to cytokines alone, particularly TNF-α and IFN-γ induces memory-like transcriptional responses, including both enhanced and dampened of gene expression upon re-stimulation. These changes resemble the priming, training, and tolerance states described in innate immune memory^19,24^, supporting the idea that ECs can encode previous inflammatory experiences in a lasting manner.

While our data reveal memory-like phenotypes, the underlying responses do not fully align with the canonical pathways or chromatin signatures of trained immunity. Genes such as *HDAC9*, *TLR2*, and *IL1B* implicated in trained immunity in monocytes.^4,25,26^ were expressed in memory-like patterns but lacked changes in chromatin accessibility in our EC ATAC-seq profiles. This suggests that, in ECs, these genes may function as stable trans-regulators rather than undergoing direct epigenetic remodeling themselves. Future studies will be needed to map their downstream regulatory networks and clarify how they contribute to EC memory-like behavior.

Notably, TNF-α and IFN-γ elicited distinct transcriptomic and chromatin responses, with limited gene overlap. TNF-α re-stimulation predominantly activated pathways linked to infection and leukocyte recruitment, while IFN-γ engaged autoimmune and interferon-related pathways. These results point to cytokine-specific imprinting mechanisms, consistent with the idea that chronic inflammation does not elicit a uniform endothelial response but one that is shaped by the nature of the inflammatory stimulus.^27^ The limited overlap of genes activated upon the repeated exposure to TNF-α and IFN-γ suggests that cytokine-specific effects contribute to the diversity of EC responses. In support of this, chromatin accessibility changes identified also showed a lack of shared chromatin regions responding similarly to TNF-α and IFN-γ. Together, these observations suggest distinct cytokine-specific effects at both the transcriptomic and epigenetics levels, adding complexity to the mechanisms underlying inflammatory memory in endothelial cells.

To validate the functional significance of these molecular changes, we used a micro-physiological relevant^28^ blood-vessel on chip model^18^ and demonstrated that TNF-α re-exposure enhances monocyte adhesion and the expression of adhesion molecules like ICAM-1 and VCAM processes central to inflammation and vascular pathology.^9^ However, we also observed inter-donor variability in HUVEC responses, with some individuals showing enhanced responses and others showing tolerance or minimal changes. This inter-individual heterogeneity aligns with prior findings on the genetic regulation of vascular inflammation and highlights the need for personalized models to study endothelial reprogramming.^29^

Beyond functional consequences, we examined how DORs in response to cytokine exposure overlapped with GWAS-SNPs linked to immune functions and diseases. These identified variants associated with COVID-19, septic shock, vaccine responses to hepatitis B and other traits related to immune function, with multiple variants located in or near MHC class II genes. While antigen presentation by MHC class II is widely attributed to the sole ability of immune cells, increasing evidence suggests that ECs can contribute under inflammatory conditions. For example, endothelial compartments in the lung and kidney express HLA-DR to shape CD4-positive memory T-cell proliferation and regulatory T-cells in inflammatory environments.^30,31^ In line with this, we observed that repeated IFN-γ exposure induced expression of all major components of MHC class II (HLA-DR, HLA-DP and HLA-DOA), both at RNA and chromatin regulation. This aligns with previous reports showing that endothelial MHC class II expression can influence immune interactions in settings such as mitigate graft rejection^32^ and rescue immune exhaustion in septic shock patients.^22^ Given the diversity of immune-related loci in the overlap, we examined which associations were statistically supported. When testing for statistical enrichment using a permutation-based approach, cardiovascular and vascular disease associated SNPs were significantly enriched within the dynamic DORs. This suggests that while inflammatory stimulation affects loci linked to a broad range of immune traits, the strongest and most consistent epigenetic associations point to vascular disease risk.

Despite the novelty of these findings, several limitations should be acknowledged. The in vitro model used for chronic inflammation stimulation, which involves repetitive exposure to TNF-α or IFN-γ, simplifies the complex in vivo environment. While these cytokines play pivotal roles in immune responses, other cytokines, and inflammatory mediators contribute to the complex environment during chronic inflammation. The duration of cytokine stimulation was relatively short, with 24 hours for the first exposure and 4 hours for the second exposure. Longer-term stimulation might elicit additional or different responses, especially considering the chronic nature of inflammation observed in certain diseases. Additionally, the resting period of 48-hours between stimulations may not fully reflect the dynamics of cytokine exposure in chronic inflammation, where inflammatory signals can persist or fluctuate over longer periods. For future studies extending this resting period may help better delineate the distinction between residual activation and true memory-like responses. Lastly, while the blood-vessel-on-chip experiments provide functional validation, they have inherent complexities. The observed inter-individual variability in monocyte adhesion underscores the need for a more comprehensive understanding of the factors influencing the memory-like responses of endothelial cells.

In conclusion, our findings contribute to the evolving understanding of endothelial cell responses to chronic vascular inflammation. These reprogrammed states are cytokine-specific, functionally significant, and genetically linked to disease-associated loci. While not mechanistically identical to trained immunity, these memory-like features represent a crucial extension of the concept to the vascular endothelium. As such, they offer a new perspective on how chronic inflammation drives long-term vascular dysfunction and point to endothelial memory as a potential therapeutic target in vascular diseases.

## Acknowledgements

The authors thank the lab members in the Genetics department, UMCG for supporting the experiments and brainstorming the findings, especially Kate McIntyre for editing English in the manuscript. This work was supported by the Netherlands Organ-on-Chip Initiative, an NWO Gravitation project (024.003.001) funded by the Ministry of Education, Culture, and Science of the government of The Netherlands (KL, HM, AvdW, SW). I.J. is supported by a Rosalind Franklin Fellowship from the University of Groningen and an NWO VIDI Grant (No. 016.171.047). VK is supported by a Hypatia tenure track Fellowship from the Radboud UMC. NK is supported by the ZonMw (10430012010002) awarded to VK.

## Author contributions

KL, VK, MN contributed to the study concept. KL, VK designed the study and experiments. AB, VO, LJ, MN, IJ, SW, AM gave feedback on the experimental design, provided facility, resources and discussed the outcomes. KL, TLD conducted cell stimulation experiments in well plates, KL collected materials and conducted RNAseq and ATACseq data. HM performed experiments on chip models and analyzed adhesion assay data. NK performed QC and analyzed RNAseq and ATACseq data. The first draft of the manuscript was written by KL and all authors gave feedback. All authors read and approved the final manuscript.

## Data availability

The RNA-seq and ATAC-seq data generated during the study have been deposited in GEO under accession number GSE267477 and GSE267418. Processed data and other relevant files are available as supplementary information. Additional data supporting the findings are available from the corresponding author upon reasonable request.

---

**Table 1:**
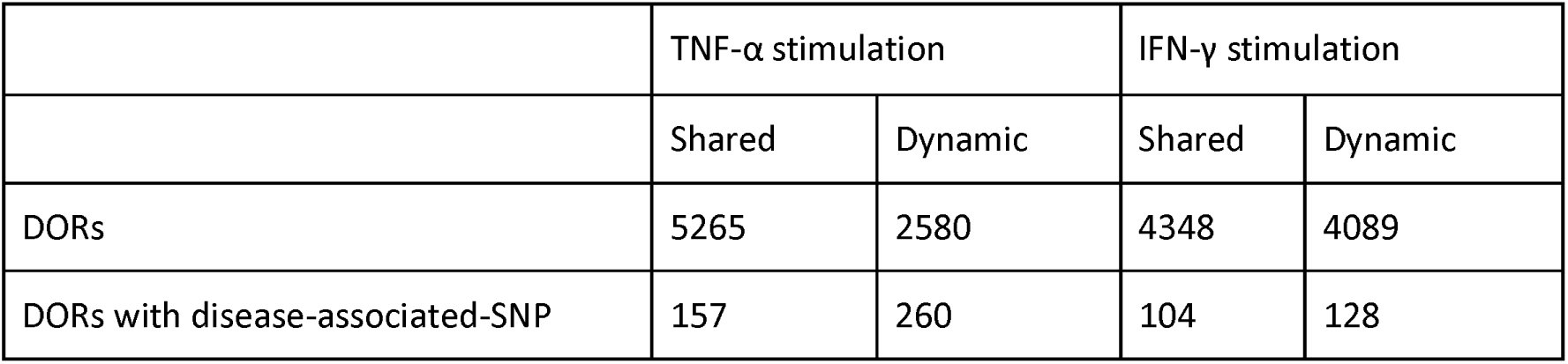
Number of ATAC-seq DE peaks and GWAS-SNPs.

| | TNF-α stimulation | | IFN-γ stimulation | |
| --- | --- | --- | --- | --- |
|  | Shared | Dynamic | Shared | Dynamic |
| DORs | 5265 | 2580 | 4348 | 4089 |
| DORs with disease-associated-SNP | 157 | 260 | 104 | 128 |

**Figure S1:**
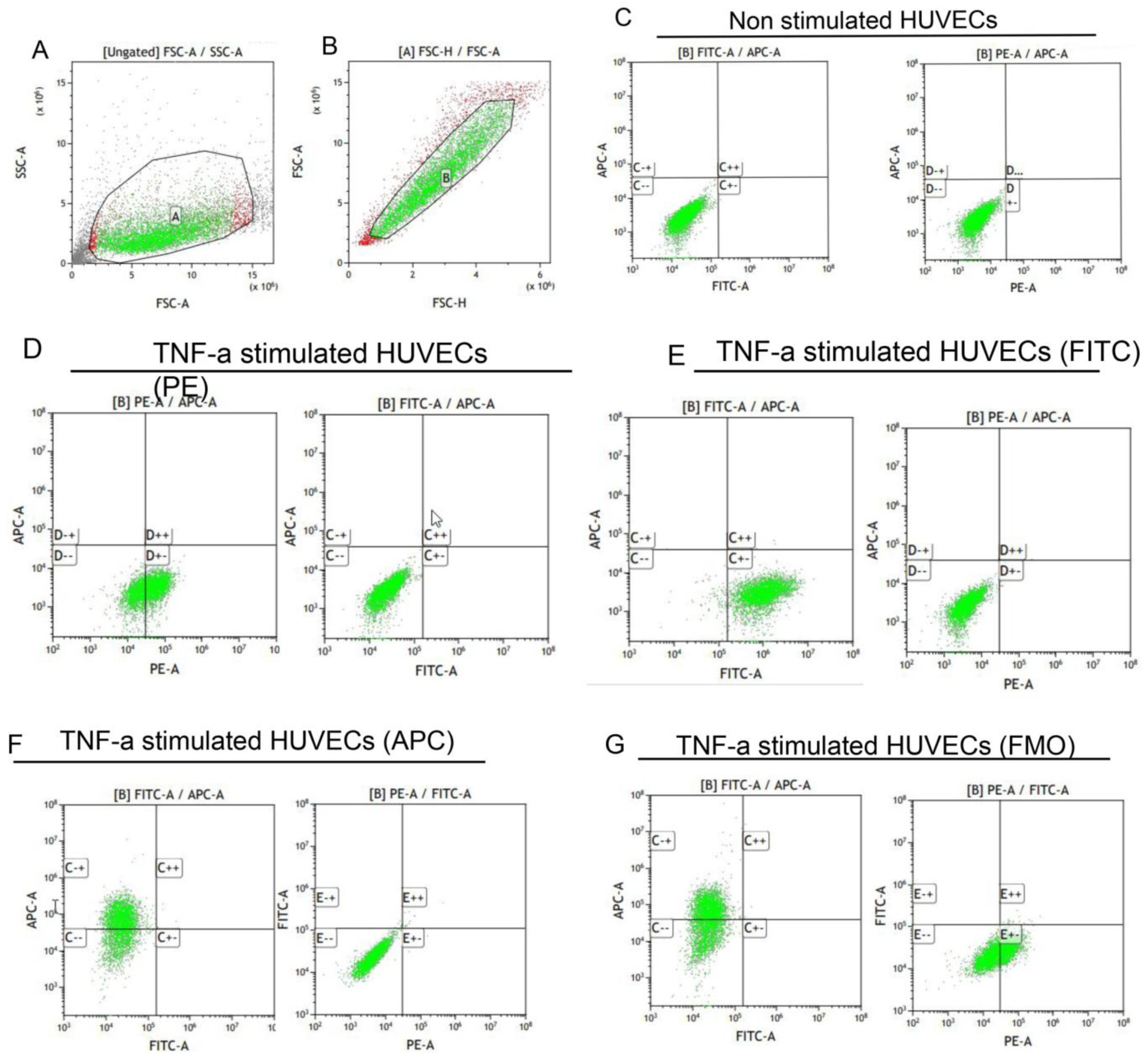
Gating strategy for flow cytometry analysis. Each dot or point on the plot is a representation of an individual cell passing through the laser. Intact cells were selected by gating an area illustrated by forward scatter (FSC-A) and side scatter (SSC-A) **(A)**. Single cells were selected based on the FSC-H and FSC-A **(B)**. Unstimulated cells, stained with IgG isotope controls were used for setting the gate for each florescent marker. Muti-color compensation was calibrated using activated HUVECs (TNF-α for 4 hours) **(C)**. E-selectin-positive cells (PE) were identified, while ICAM-1 (APC) and VCAM-1 (FITC) signals remained negative **(D)**, ICAM-1 positive cells (FITC) were identified, while E-selectin (PE) and VCAM-1 (APC) remained negative. **(E).** VCAM-1 positive cells (APC) were identified while E-selectin (PE) and ICAM (FITC) remained negative **(F)**. A fluorescence minus one (FMO) control was included, where all markers were stained, except FITC **(G).**

**Figure S2:**
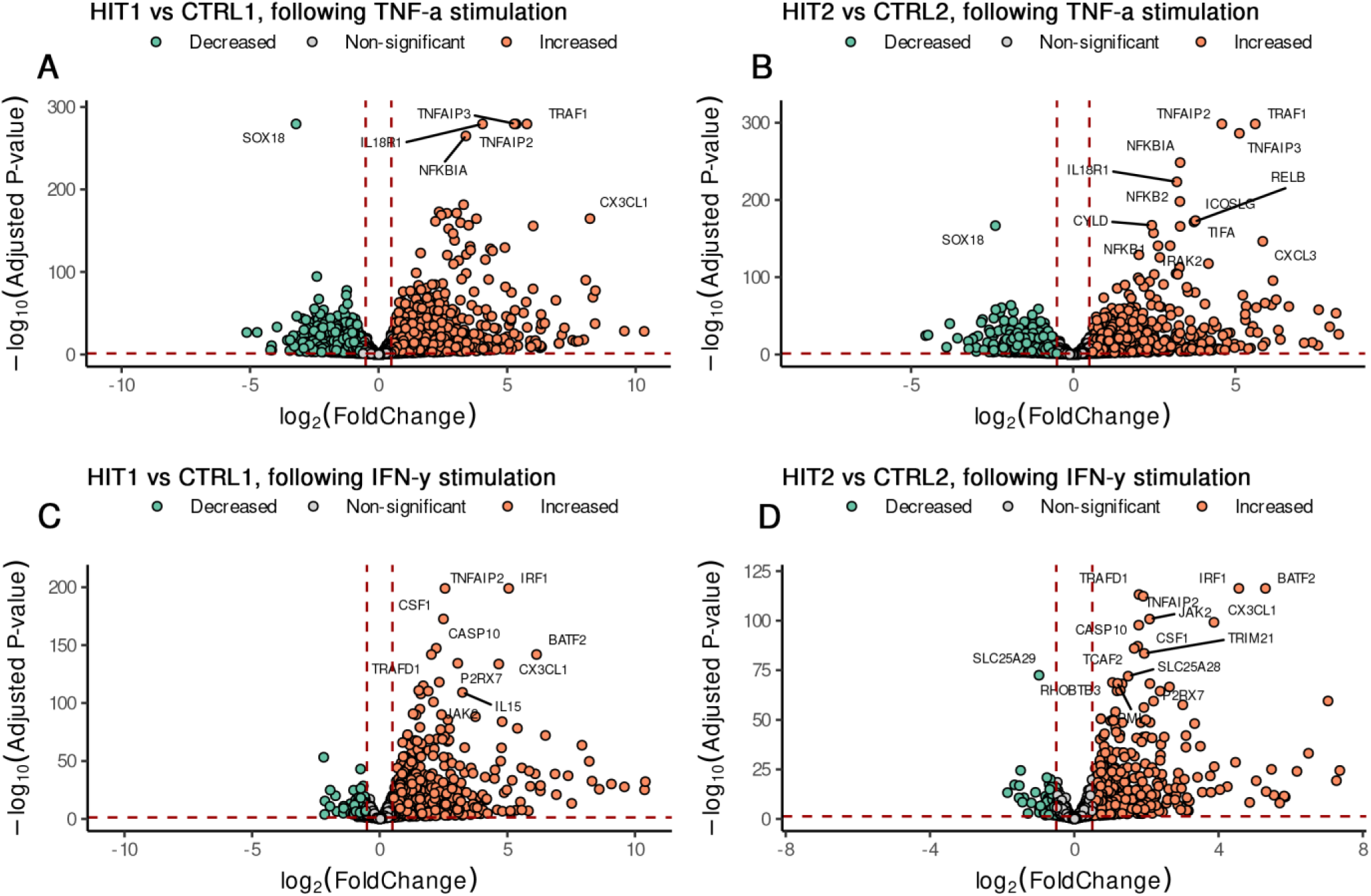
Volcano plots depicting transcriptional changes in ECs upon TNF-α and IFN-γ stimulation. Volcano plots showing DEGs identified in different conditions, HIT1 vs CTRL1 **(A)** and HIT2 vs CTRL2 **(B)** for TNF-α stimulation, and HIT1 vs CTRL1 **(C)** and HIT2 vs CTRL2 **(D)** for IFN-γ stimulation. Increased (orange) and decreased (green) DEGs are highlighted, with top 10 genes labeled. DEGs were filtered based on FDR adjusted p-values ≤ 0.05 and log2 fold change (log2FC ≥ 0.5) thresholds. The x-axis represents the log₂ fold change, while the y-axis visualizes the -log₁₀ transformed FDR-adjusted p-values.

**Figure S3:**
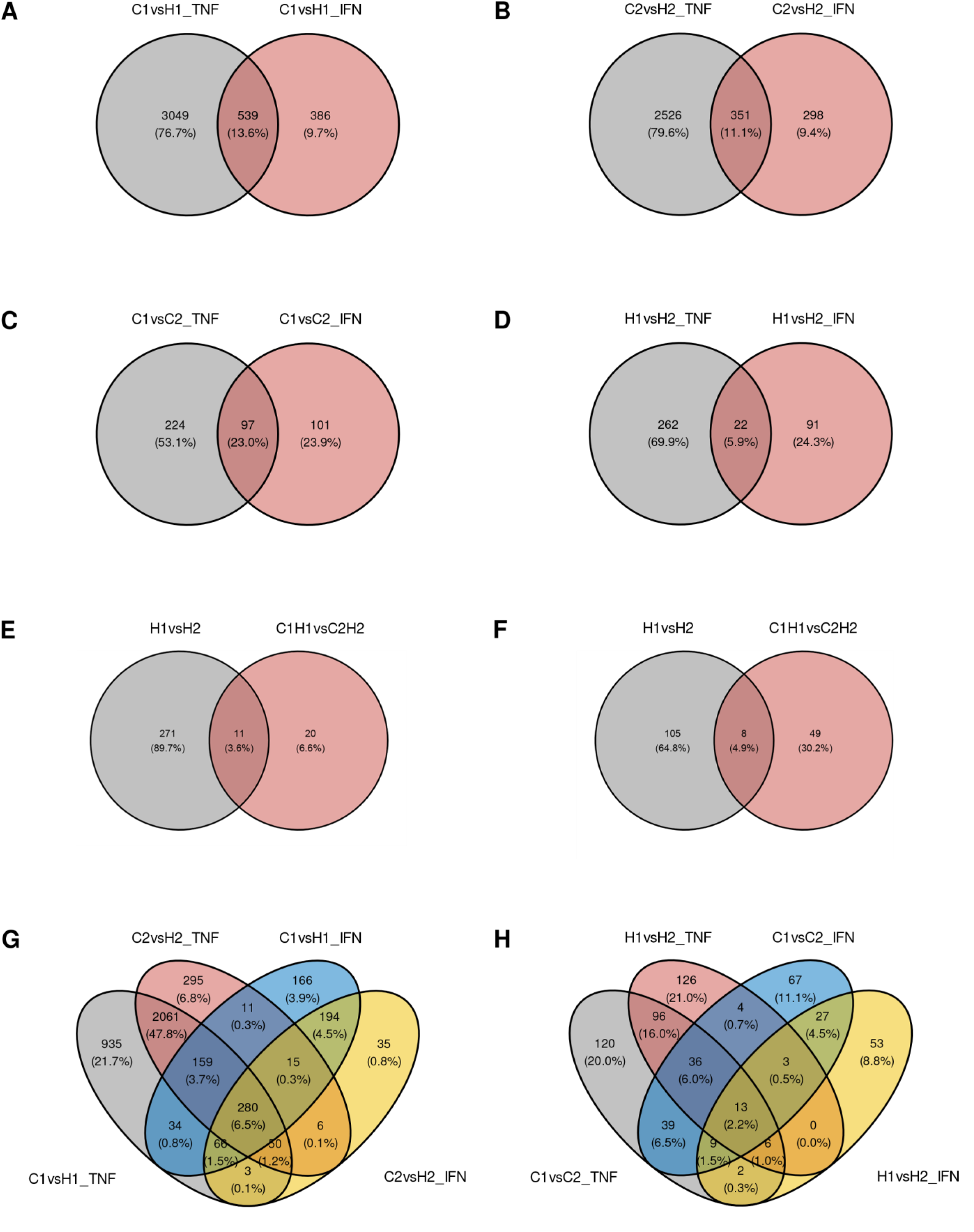
Overlapping genes between different stimulation conditions. Venn diagram visualizing the overlapping genes between the TNF-a and IFN-y stimulation after single stimulation (HIT1 vs CTRL1) **(A)** and repeated stimulation (HIT2 vs CTRL2) stimulation **(B)**. Whereas the overlapping genes between the control conditions (CTRL2 vs CTRL1) between TNF-a **(C)** and IFN-y **(D).** Overlapping genes between the stimulation (HIT2 vs HIT1) and (HIT2 – CTRL2 vs HIT1 – CTRL1) conditions are depicted for TNF-α **(E)** and IFN-y **(F).** Lastly, we compared the first (HIT1 vs CTRL1) and second (HIT2 vs CTRL2) stimulation of both cytokines **(G),** and the control (CTRL2 vs CTRL1) and stimulated (HIT2 vs HIT1) conditions **(H).**

**Figure S4:**
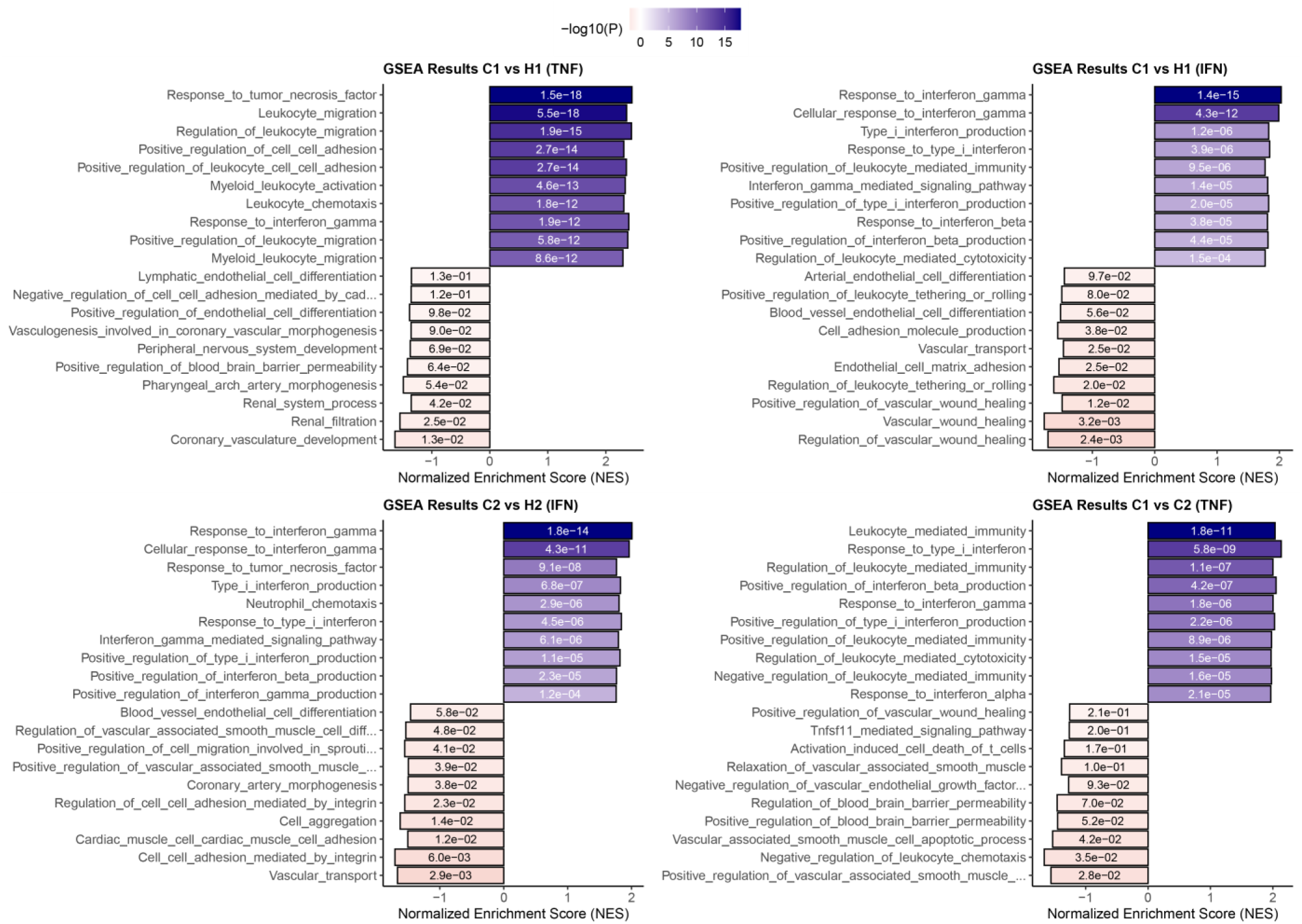
Enrichment of endothelial specific pathways across conditions. Gene set enrichment analysis (GSEA) was performed using curated endothelial specific gene sets derived from GO biological process. Results for the single-hit stimulation (HIT1 vs CTRL1) **(A)** and repeated expose (HIT2 vs CTRL2) **(B**) following TNF-α stimulation. Similarly, results of the single-hit stimulation (HIT1 vs CTRL1) **(C)** and repeated exposure (HIT2 vs CTRL2) **(D**) following IFN-γ stimulation. Blue bars indicate upregulated pathways, while red bars indicate downregulated pathways. The x-axis indicates the normalized enrichment score (NES), and the intensity of the colors reflects the –log10 FDR adjusted p-value

**Figure S5:**
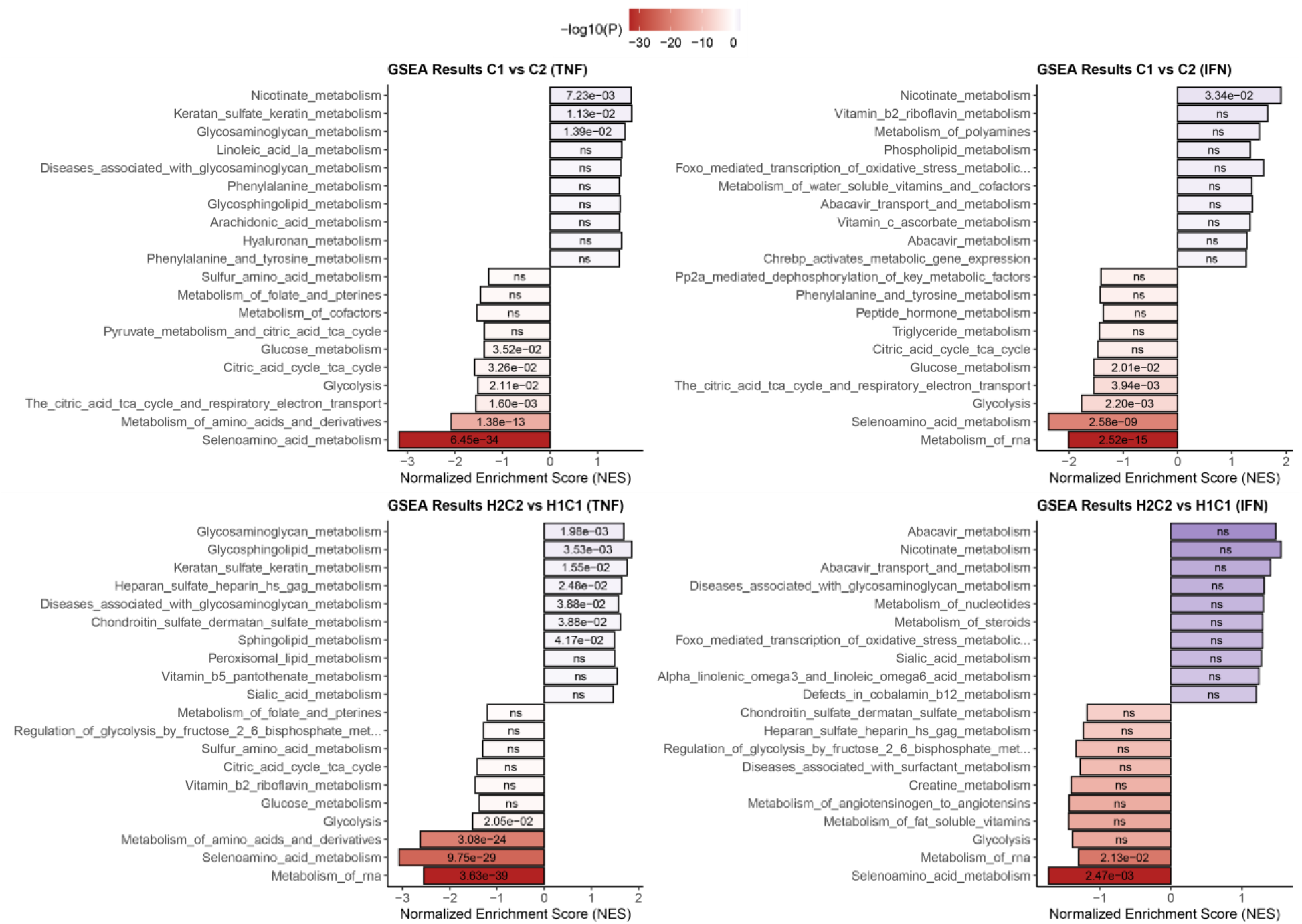
Enrichment of metabolic specific pathways across conditions. Gene set enrichment analysis (GSEA) was performed using metabolic specific gene sets derived from Reactome. Results for the control conditions (CTRL2 vs CTRL1) **(A)** and baseline-adjusted comparison(HIT2 - CTRL2 vs HIT1 – CTRL1) **(C**) following TNF-α stimulation. Similarly, results of the control stimulation (CTRL2 vs CTRL1) **(B)** and baseline adjusted (HIT2 - CTRL2 vs HIT1 – CTRL1) **(D**) following IFN-γ stimulation. Blue bars indicate upregulated pathways, while red bars indicate downregulated pathways. The x-axis indicates the normalized enrichment score (NES), and the intensity of the colors reflects the –log10 FDR adjusted p-value

**Figure S6:**
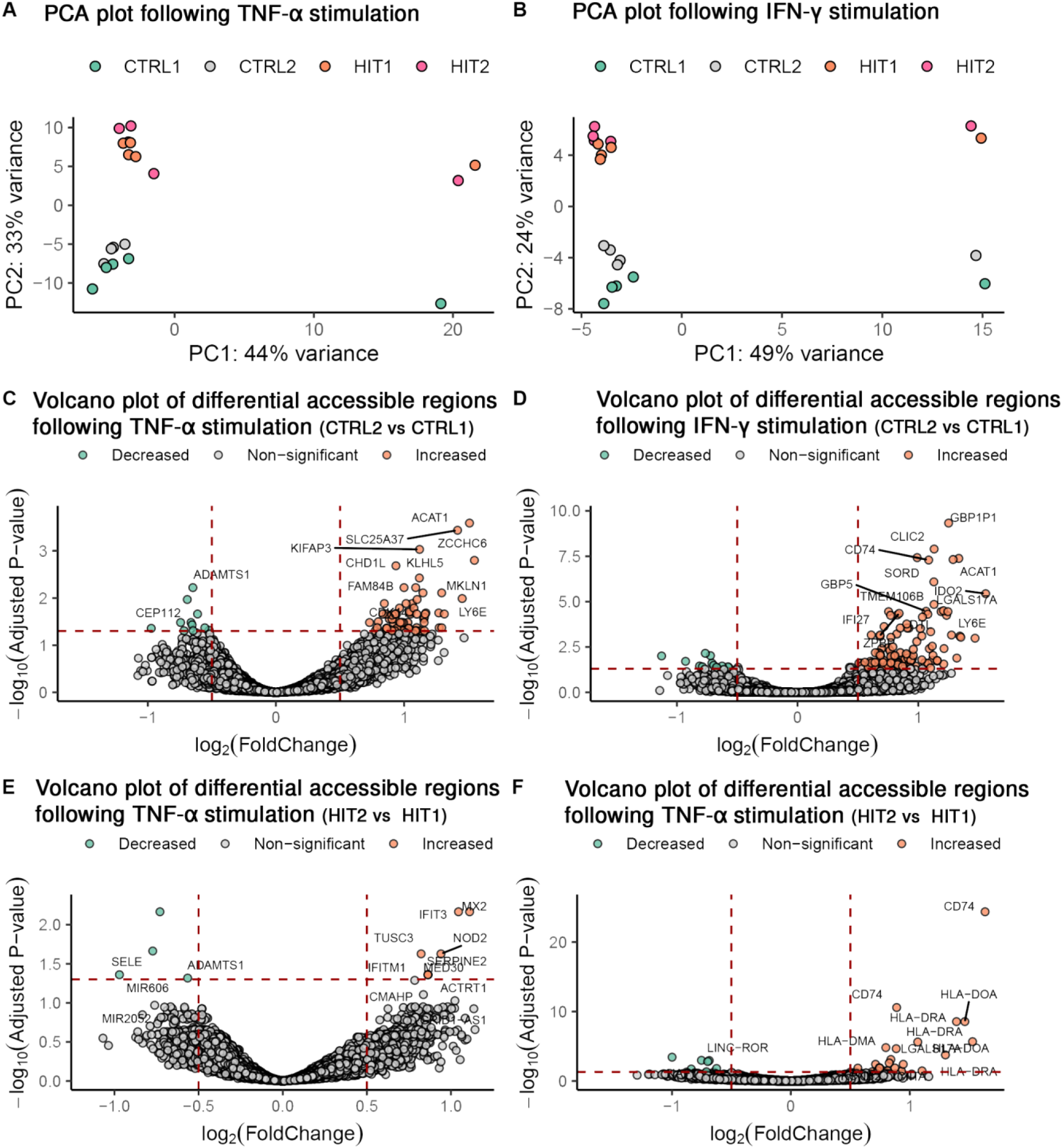
PCA and volcano plots depicting chromatin accessibility and transcriptional changes in ECs upon TNF-α and IFN-γ stimulation. PCA plots show the variance in chromatin accessibility across four conditions following TNF-α stimulation **(A)** and IFN-γ stimulation **(B).** Colors depict the different stimulation conditions. Volcano plots showing DORs identified in different conditions, CTRL2 vs CTRL1 for TNF-α **(B)** and IFN-γ **(C),** HIT2 vs HIT1 for TNF-α **(D)** and IFN-γ. Increased (orange) and decreased (green) DORs are highlighted, with top 10 DORs are labeled with gene names. DORs were filtered based on FDR adjusted p-values ≤ 0.05 and log2 fold change thresholds.

**Figure S7:**
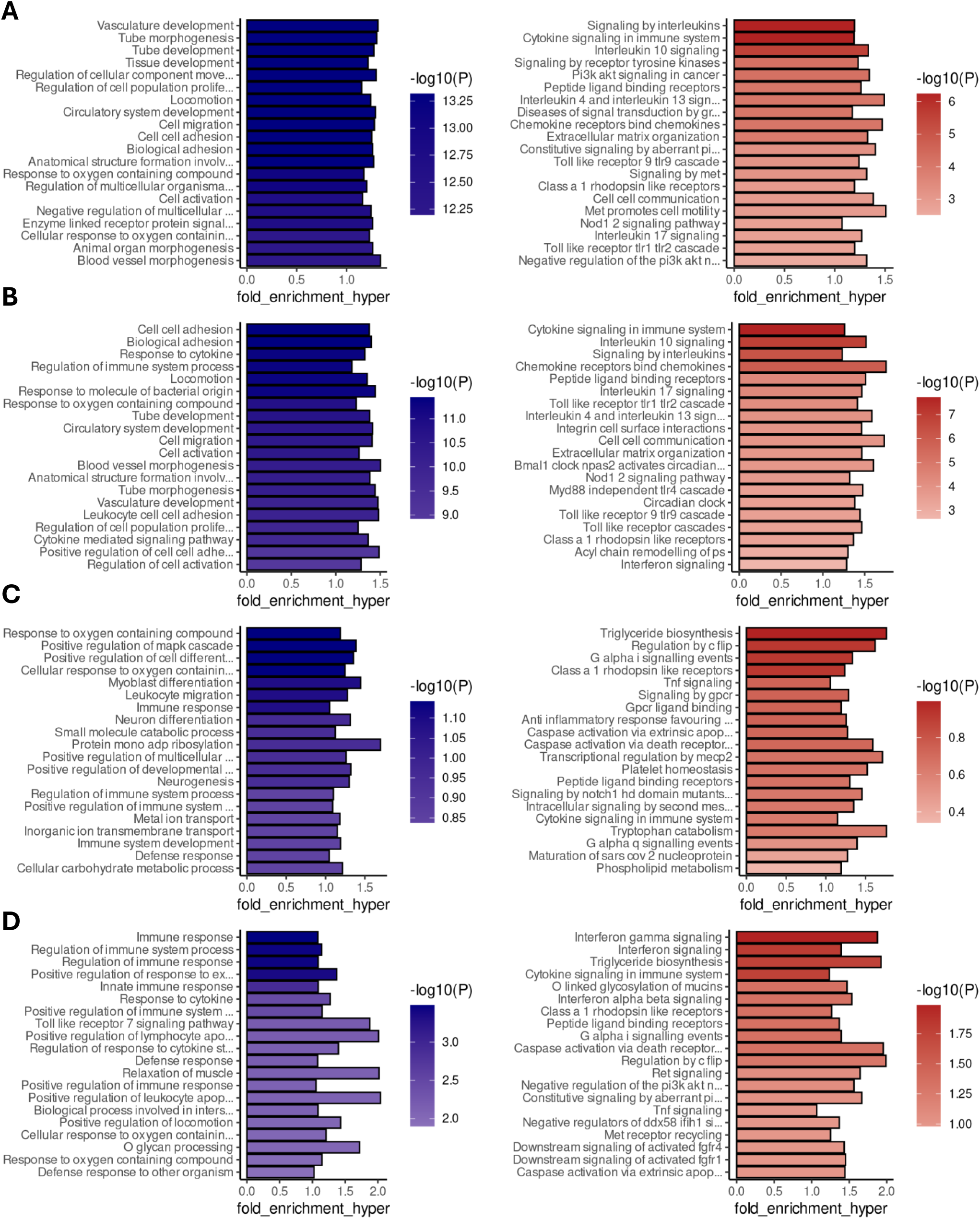
Enrichment results of DORs across conditions using both GO biological process and Reactome. Pathway enrichment results for the single-hit stimulation (HIT1 vs CTRL1) **(A)** and repeated expose (HIT2 vs CTRL2) **(B**) following TNF-a stimulation. Similarly, results of the single -hit stimulation (HIT1 vs CTRL1) **(C)** and repeated expose (HIT2 vs CTRL2) **(D**) following IFN-y stimulation. Blue bars represent results for GO biological process, while red bars represents Reactome pathways. The x-axis indicates the enrichment score, while the intensity of the colors represents the –log10 FDR adjusted p-value

**Figure S8:**
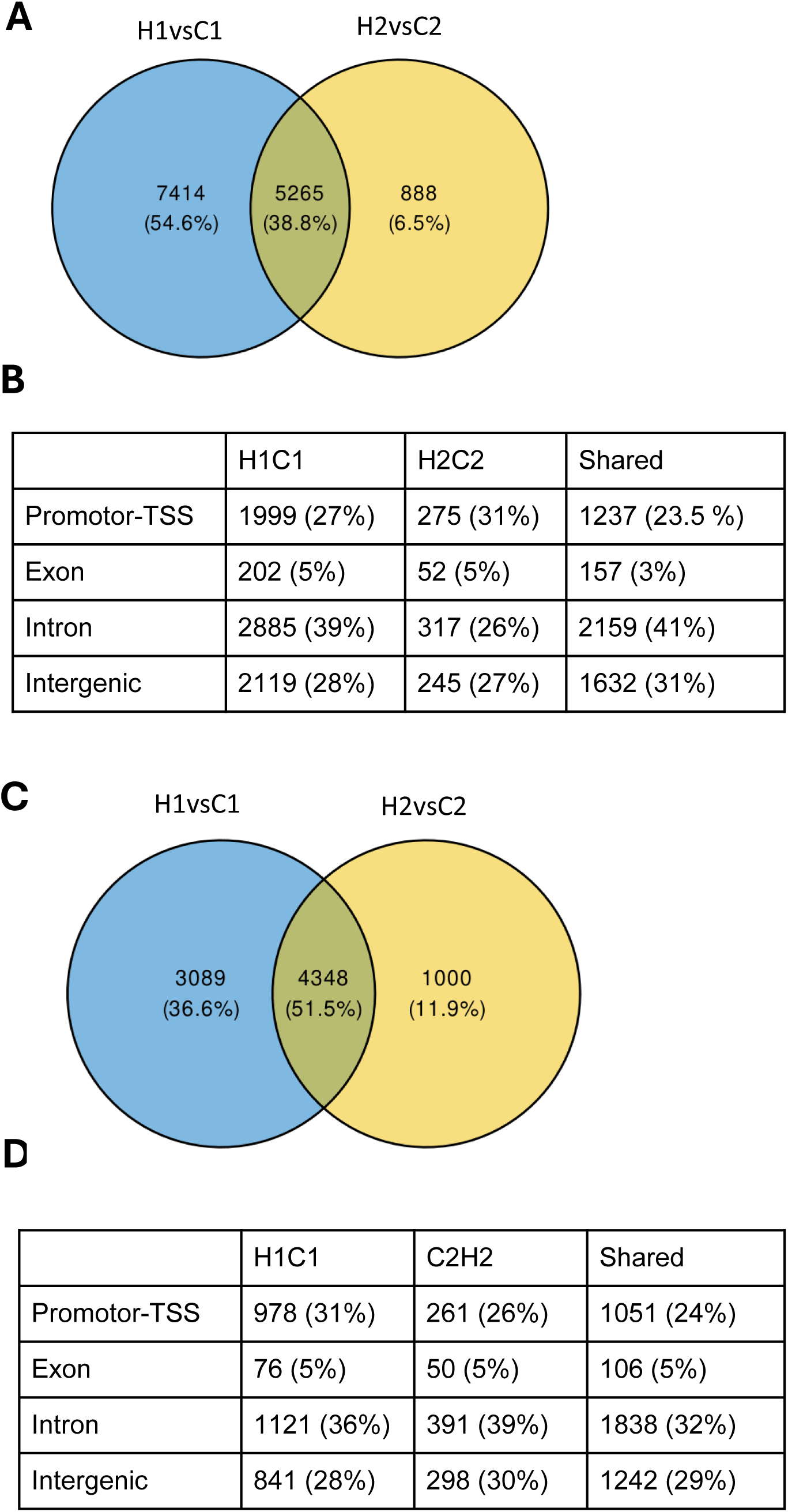
Overlap and genomic distribution of differentially open regions (DORs) upon single and repeated cytokine stimulation. Venn diagrams visualizing the overlapping DORs between the single stimulation (HIT1 vs CTRL1) and repeated stimulation (HIT2 vs CTRL2) for TNF-α **(A),** and IFN-γ **(C)**. Percentages represent the proportion of DORs unique to each condition or common between them. Distribution of different genomic regulatory regions are of DORs for TNF-α **(B)** and IFN-γ **(D).** The table provides the total number and the percentages of DORs in each category.

**Figure S9:**
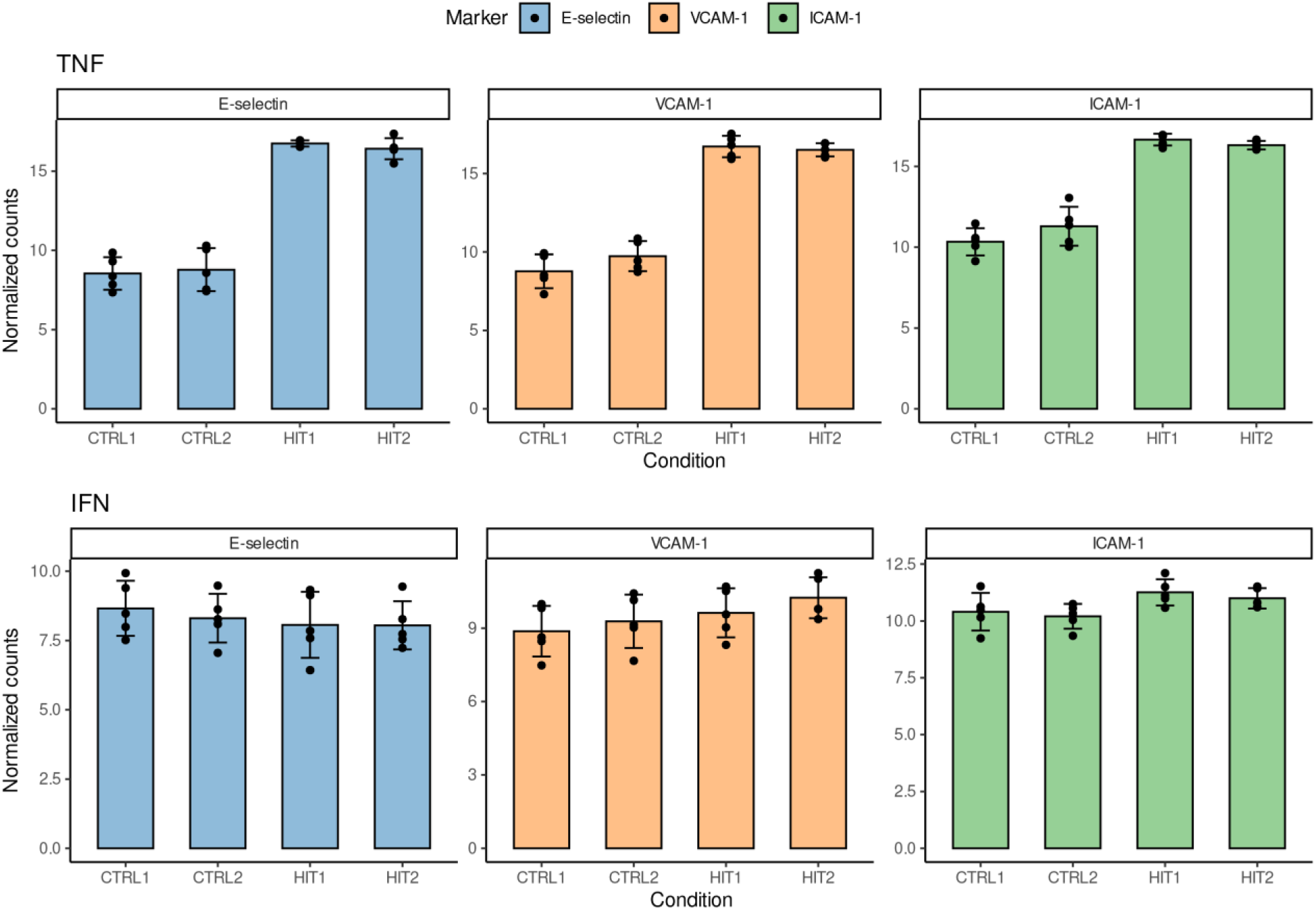
Transcriptional profiles of adhesin molecules. Bar plot representing the RNA-seq transcript levels of adhesion molecules E-selectin, VCAM-1, and ICAM-1 under different experimental conditions, following cytokine stimulations. Error bars indicate the standard error of the mean (SEM) derived from biological replicates (n = 5). The y-axis represents the normalized RNA-seq count data, while the x-axis represent different stimulation conditions.

